# Ageing-associated myelin dysfunction drives amyloid deposition in mouse models of Alzheimer’s disease

**DOI:** 10.1101/2021.07.31.454562

**Authors:** Constanze Depp, Ting Sun, Andrew Octavian Sasmita, Lena Spieth, Stefan A. Berghoff, Agnes A. Steixner-Kumar, Swati Subramanian, Wiebke Möbius, Sandra Göbbels, Gesine Saher, Silvia Zampar, Oliver Wirths, Maik Thalmann, Takashi Saito, Takaomi Saido, Dilja Krueger-Burg, Riki Kawaguchi, Michael Willem, Christian Haass, Daniel Geschwind, Hannelore Ehrenreich, Ruth Stassart, Klaus-Armin Nave

**Author notes:** **Correspondence** Correspondence and requests for materials, data or code should be addressed to or. shared first authorship (Constanze Depp, Ting Sun).

## Abstract

The prevalence of Alzheimer’s disease (AD), the leading cause of dementia, shows a strict age-dependency, but why ageing constitutes the main risk factor for this disease is still poorly understood. Brain ageing affects oligodendrocytes^1^ and the structural integrity of myelin sheaths^2^, the latter associated with secondary neuroinflammation^3^. Since oligodendrocytes support axonal and neuronal health^4–7^, we hypothesised that ageing-associated loss of myelin integrity could be an upstream risk factor for neuronal amyloid-β (Aβ) deposition, the primary neuropathological hallmark of AD. Here, we show that in AD mouse models different genetically induced defects of myelin integrity or demyelinating injuries are indeed potent drivers of amyloid deposition *in vivo*, quantified by whole brain light sheet microscopy. Conversely, the lack of myelin in the forebrain provides protection against plaque deposition. Mechanistically, we find that myelin dysfunction causes the accumulation of the Aβ producing machinery within axonal swellings and increases cortical amyloid precursor protein (APP) cleavage. Surprisingly, AD mice with dysfunctional myelin lack plaque-corralling microglia but show a disease-associated microglia (DAM)-like signature as revealed by bulk and single cell transcriptomics. These activated microglia, however, are primarily engaged with myelin, preventing the protective reactions of microglia to Aβ plaques. Our data suggest a working model, in which age-dependent structural defects of myelin promote plaque formation, directly and indirectly, and are thus an upstream AD risk factor. Improving oligodendrocyte health and myelin integrity could be a promising target to delay AD.g

## Main text

The pathology of AD is characterised by the deposition of Aβ plaques and neurofibrillary tangles primarily in cortex and hippocampus^8^. According to the ‘amyloid hypothesis’ of AD, the build-up of Aβ initiates a cascade of harmful events that lead to neuronal dysfunction^9^. More than lifestyle choices and genetic predisposition, old age is the primary risk factor for AD development, but exactly how brain ageing is linked to amyloid deposition is unclear^10^. Myelin, a spirally wrapped and compacted glial membrane, enhances axonal conduction speed^11^ and its non-compacted regions allow oligodendrocytes to provide metabolic support to the neuronal compartment^5,6,12^. The unique cellular architecture of myelin makes protein and lipid turnover challenging and slow^13–15^. This, together with the long lifetime of oligodendrocytes^16^ and age-dependent loss of epigenetic marks, might explain the structural deterioration of myelin with age^2^. We speculated that the breakdown of myelin integrity in the ageing brain acts as driving force for Aβ deposition and tested this hypothesis in mouse models of amyloidosis, in which we introduced myelin dysfunction of various degrees by genetic and pharmacological manipulations.

### Intracortical myelin decline in AD patients and mouse models

Macroscopic brain imaging studies have suggested that cortical myelin damage occurs in the preclinical phase of AD, i.e., prior to extensive amyloid deposition^17–21^. Microscopic evidence of this, however, remains scarce. We therefore first studied intracortical myelin integrity in a small cohort of AD patients, focussing on the trans-entorhinal area by immunofluorescence analysis of myelin (Fig.1a). We observed a striking decline of intracortical myelin density in autopsies from AD patients which were not limited to the immediate vicinity of plaques. Sites of myelin loss were also associated with increased numbers of Iba1^+^ microglia. Since human neuropathological correlations cannot distinguish between myelin loss being the cause or consequence of neuronal AD pathology such as axonal decay, we turned to two different experimental animal models of AD (5×FAD, APP^NLGF^). Here, we are in a position to ask whether loss of myelin can act as an upstream driver of amyloidosis by combining AD mouse models with mice developing subtle genetically-induced myelin disintegration (Fig.1b). Specifically, we employed null mutants of the myelin architectural proteins CNP and PLP1, that display rather minor structural myelin defects. Lack of CNP causes a collapse of cytosolic channels within the myelin sheath^22^ associated with axonal swellings^23^. PLP1-deficient myelin lacks physical stability^24^, reduces axonal energy metabolism^25^ and causes axonopathy at higher age^4^. We first compared normal brain ageing of c57Bl6 wildtype mice with that of *Cnp* and *Plp1* null mutants on the same background. By immunostaining, *Cnp* and *Plp1* mice showed gliosis in grey and white matter at 6 months, qualitatively similar to aged wildtype mice (24-month-old), but more profound (Fig.1c). Similar to AD patients, aged wildtype mice and more pronounced 6-month-old myelin mutants displayed significant reductions of intracortical myelin content, especially in the upper cortical layers (Fig.1c).

**Figure 1.**
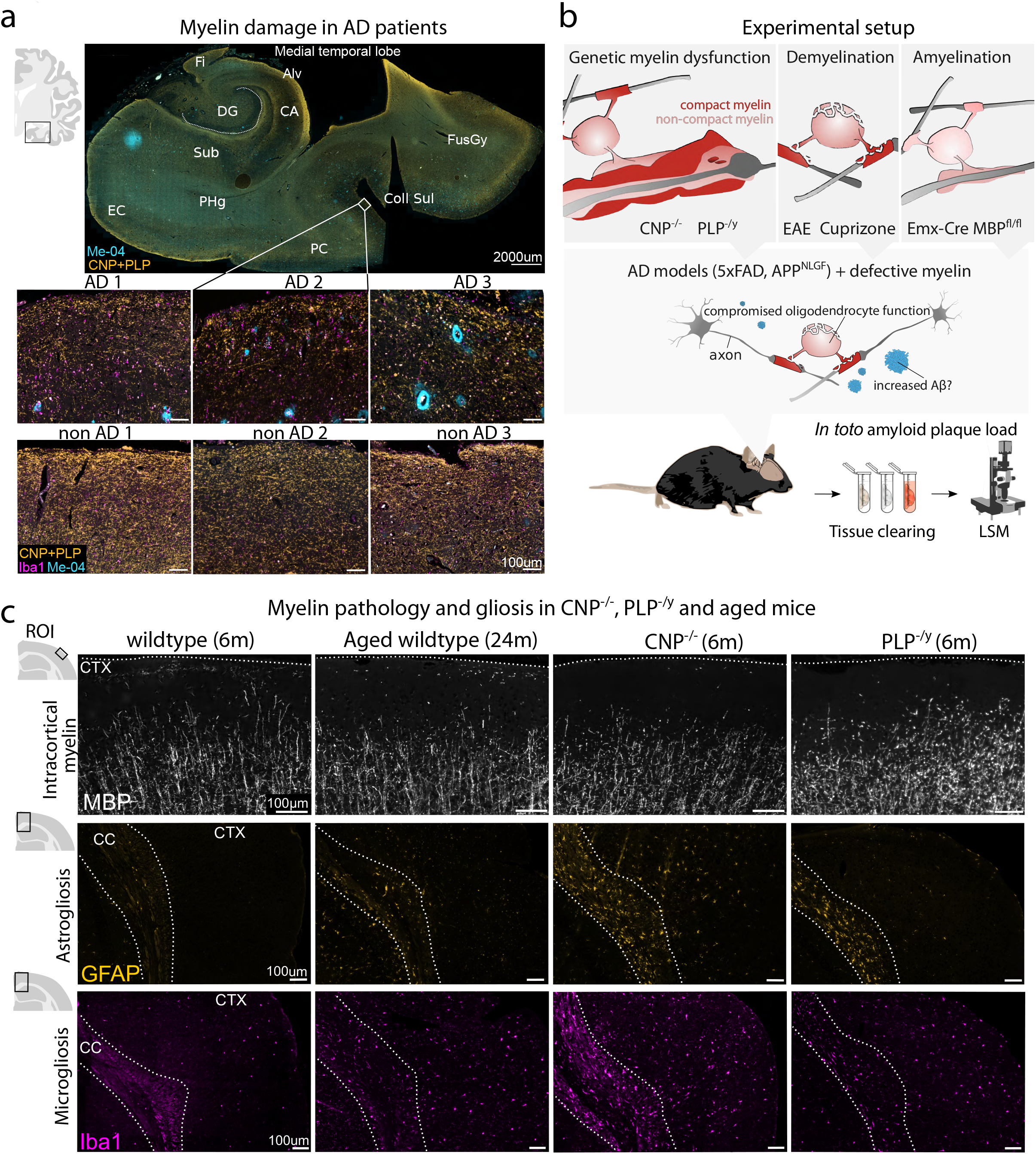
Myelin mutant animals show premature myelin integrity breakdown recapitulating myelin decline in ageing and AD. **(a)** Myelin damage in Alzheimer’s disease (AD) patients in the medial temporal lobe visualised by immunolabelling against CNP and PLP1 to stain myelinated fibres, Iba1 to stain microglia and Methoxy-04 (Me-04) to stain amyloid-β plaques. Annotated overview image of the medial temporal lobe of one AD patient indicates the location of the closeups, which show upper cortical layers in the transentorhinal cortex in plaque^+^ individual AD patients and non-AD controls (n=3 per group). DG: Dentate Gyrus, Fi: Fimbria, CA: Cornu Ammonis, Sub: Subiculum, EC: Entorhinal cortex. PHg: Parahippocampal gyrus. FusGy: Fusiform gyrus. Coll Sul: Collateral Sulcus. **(b)** Experimental setup to study the effect of myelin dysfunction on amyloid plaque load in AD mouse models. Models of genetic myelin dysfunction, acute demyelination and genetic amyelination were combined with the 5×FAD and APP^NLGF^ model of AD and effects on amyloid deposition were investigated in toto by tissue clearing and light sheet microscopy (LSM). **(c)** Myelin defects drive a premature ageing phenotype in mice. Panels show microscopic images of immunolabellings against MBP (Myelin), GFAP (Astrogliosis), IBA1 (Microgliosis) in 6-month-old wildtype, 24-month-old wildtype, 6-month-old CNP^-/-^ and 6-month-old PLP^-/y^ mice. Regions of interest (ROI) analysed are indicated on the left. Dashed lines in the upper panel marks pial surface. Dashed line in lower panels outline the corpus callosum. CC: Corpus callosum. CTX: Cortex.

### Myelin dysfunction drives amyloid deposition in AD mouse models

To determine whether such ageing-associated myelin defects can drive amyloid deposition, we crossbred *Cnp*^*-/-*^ and *Plp1*^-/*y*^ mutants with commonly used mouse models of AD and analysed plaque burden in the resulting offspring (Fig.1b). For this, we optimised a Congo Red *in toto* plaque staining and clearing protocol based on the iDISCO technique^26,27^ for light sheet microscopy (Fig.1b, Extended data Fig.1) to determine the amyloid plaque load of the entire brain in an unbiased fashion. Indeed, when compared to 5xFAD mice, both *Cnp*^*-/-*^ *5×FAD* and *Plp1*^-/*y*^ *5×FAD* double mutants showed striking increases of amyloid plaque load, as quantified in hippocampal white matter (alveus) and cortex at 6 months (Fig.2a,b). In both models, the effects were expectedly strongest in the alveus, where Aβ was deposited in very small aggregates, indicating an increased formation of amyloid seeds. To exclude that these effects are specific to the 5×FAD model and APP overexpression, we validated our findings in crossbreedings of *Cnp*^*-/-*^ and the *APP*^*NLGF*^ knock-in model, in which the humanised triple-mutant APP is expressed under control of the endogenous *App* locus (Extended data Fig.2). In both AD models, the increase in plaque load was observed at 6 months, but not yet at 3 months when myelin defects and reactive gliosis are only subtle in *Cnp*^*-/-*^ and *Plp1*^*-/y*^ mice (Extended data Fig.3a,b). We also investigated whether myelin dysfunction modifies 5×FAD behavioural deficits by performing behavioural testing in the Y-maze (YM) and the elevated plus maze (EPM) (Extended data Fig.4). In both paradigms, double mutant mice presented with hyperactivity that was less pronounced in single mutants. In the EPM, *Cnp*^*-/-*^ *5×FAD* mice preferred the open arms, indicative of an abnormal lack of anxiety. Statistical analysis confirmed supra-additive effects (interactions) of 5×FAD and myelin mutant genotypes in many instances, also detectable in the hindlimb clasping test (Extended data Fig.4c,d). Myelin defects and amyloid pathology appear to synergistically worsen behavioural deficits indicative of disinhibition – a neuropsychiatric symptom also present in AD patients^28,29^.

**Figure 2.**
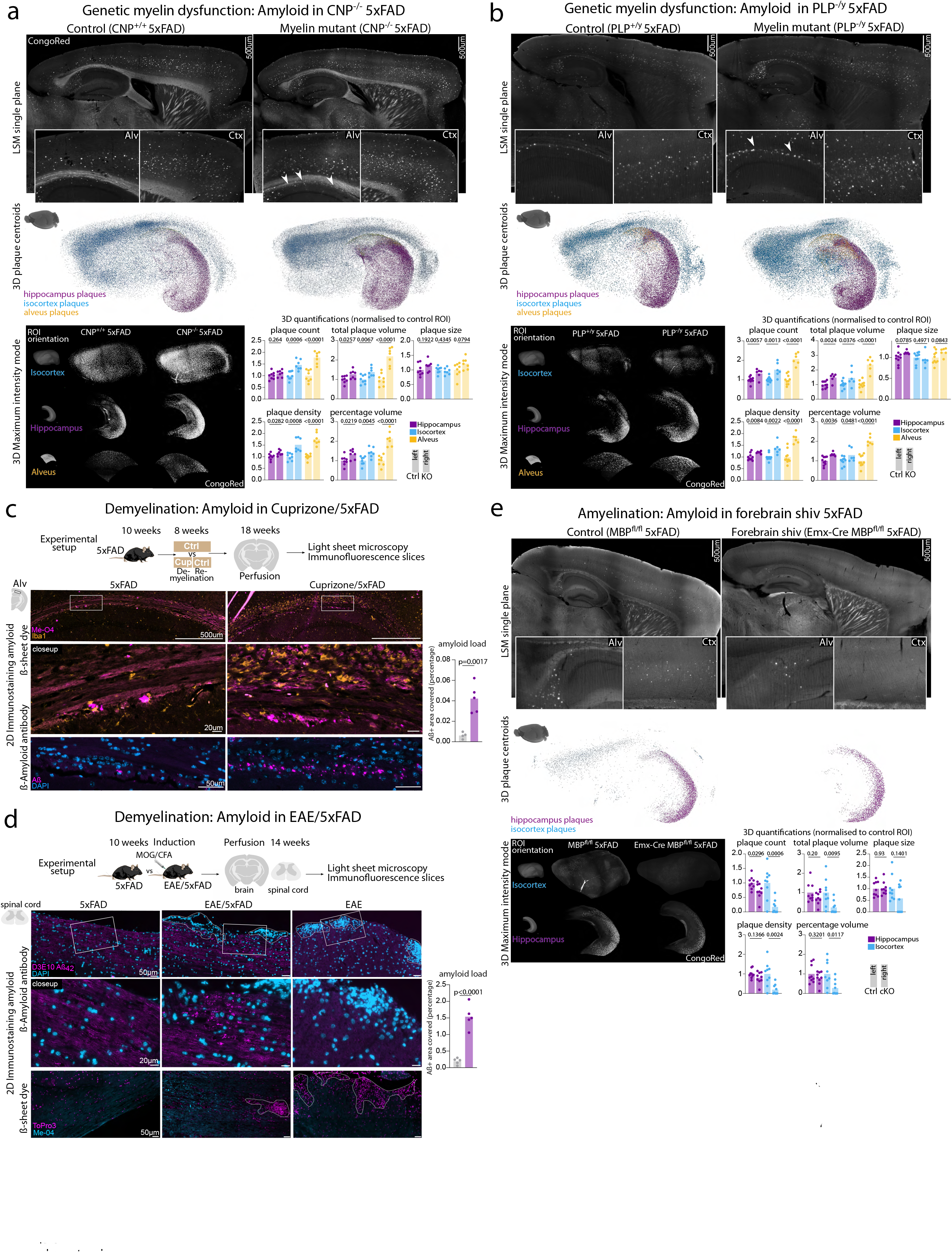
Light sheet microscopic (LSM) analysis of amyloid plaque load in genetic mouse models of dys-, de- and amyelination crossbred with 5×FAD mice. Upper panels in **(a)**, **(b)**, and **(e)** show LSM single planes of cleared brains stained for amyloid plaques by Congo Red. Inlays show closeup images of cortex and alveus. Arrows indicate small amyloid deposits. Middle panel shows 3D representation of hippocampal, isocortical and alveus plaques represented as coloured centroids. Left lower panels show 3D cropped regions of interest in rendered in maximum intensity modus. Right lower panels show the 3D quantification of plaque load in the three regions of interest normalised to the respective region in control animals. For statistical analysis of LSM data, two-sided unpaired Student’s t-test was performed. P-values are indicated above bar graphs. Dots represent single animals and bars represent mean. **(a)** In toto plaque load in 6-month-old CNP^-/-^5×FAD. n=8 for control, n=7 for mutant. **(b)** In toto plaque load in 6-month-old PLP^-/y^ 5×FAD mice. n=10 for control, n=6 for mutant. **(c)** Experimental setup to study effects of cuprizone-mediated demyelination on plaque deposition in 5×FAD mice. Middle panel shows 2D immunostaining against microglia (Iba1) and amyloid-β using the β-sheet dye Me-04 in the alveus of cuprizone-treated 5×FAD and control animals. Lower panel shows 2D Immunostainings against amyloid and quantification of amyloid positive deposits in the alveus. Dots represent single animals and bars represent mean. Statistical analysis was performed using unpaired, two-sided Student’s t-test. n=4 for control, n=5 for cuprizone treatment. **(d)** Experimental setup to study the effect of EAE-mediated demyelination on amyloid plaque load in 5×FAD animals. Lower panel shows 2D immunostainings against amyloid. EAE lesions are indicated by nuclei accumulations and marked by dashed lines. Lower panel shows β-sheet dye (Me-04) staining against amyloid plaques in EAE/5×FAD spinal cord. EAE control animals are shown to rule out unspecific staining of lesion sites in EAE. Quantification of amyloid positive deposits in the lesion environment is shown on the left. Dots represent single animals and bars represent mean. Statistical analysis was performed using unpaired, two-sided Student’s t-test. n=5 for control, n=5 for EAE treatment. **(e)** In toto plaque load in 3-month-old forebrain *shiverer* 5×FAD (Emx-Cre MBP^fl/fl^ 5×FAD). n=9 for control, n=9 for mutant.

### Acute demyelination drives amyloid deposition

To confirm that myelin dysfunction is an upstream driver of plaque pathology, we tested the effect of acute demyelination. Young adult 5×FAD mice were fed a cuprizone containing diet for 4 weeks, followed by a 4-week recovery period and the determination of plaque load via light sheet microscopy (Fig.2c). Interpreting this experiment is complicated, however, because the copper-chelating properties of cuprizone interfere with plaque core formation, which is copper-dependent^30^. Indeed, in *in toto* light sheet microscopy cuprizone treatment seemingly ameliorated hippocampal and cortical Aβ pathology (Extended data Fig.3c). However, the amyloidosis-driving effect of demyelination prevailed over the inhibition by copper-chelation in most markedly demyelinated areas such as the hippocampal alveus. Here, less-compacted Aß plaques in cuprizone treated 5×FAD could be well stained with anti-Aß antibodies revealing a strong increase in small amyloid aggregates (Fig.2c).

Independent support of our working model was provided by AD mice challenged with experimental autoimmune encephalomyelitis (EAE). We immunised young 5×FAD animals with MOG peptide and analysed their brains and spinal cord 4 weeks later (Fig.2d). Unlike an earlier study with aged J20 and Tg2576 AD mice in which EAE reportedly reduced plaque load^31^, we found no such difference in amyloid deposition in younger 5×FAD-EAE mice (Extended data Fig.3d,e). However, in the spinal cord, where demyelinating EAE pathology massively unfolds, we identified small, atypical amyloid aggregates in the peri-lesion environment, which were absent from control 5×FAD mice (Fig.2d). We verified the presence of aggregated amyloid by staining spinal cord sections with the β-sheet dye Methoxy-04 (Me-04). Of note, in EAE spinal cord from wildtype animals, no such Me-04^+^ positive material was found, ruling out the non-specific detection of lipid deposits in demyelinated lesions (Fig.2d). Taken together, myelin defects – both chronic and acute – drive amyloid deposition in AD mouse models, which identifies dysfunctional myelin as an upstream risk factor for amyloid deposition.

### Lack of myelin ameliorates amyloidosis in AD mouse models

Primate brains have more CNS myelin than smaller mammals^32^ and, compared to other apes, humans show a disproportional enlargement of prefrontal white matter^33^. This raises the question whether the extent of cortical myelination *per se* could play a causal role in human AD. We, therefore, next asked the opposite question: what impact does the near complete absence of cortical myelin have on the course of amyloidosis in 5×FAD mice? To this end, we generated a line of forebrain-specific *shiverer* mice (*Emx-Cre Mbp^flox/flox^*), in which cortical axons are largely unmyelinated, and crossbred them to 5×FAD mice (Extended data Fig.5a). At 3 months of age, forebrain *shiverer* 5×FAD mice were strongly protected against amyloid deposition, both in hippocampus and cortex (Fig.2e). However, at 6 months this effect was largely lost, revealing a delay of plaque formation in the absence of myelin (Extended data Fig.5b). We conclude that myelin ensheathment and even more so defective myelin is an upstream driving force of neuronal plaque deposition.

### Myelin dysfunction shifts neuronal APP metabolism

How do myelin defects mechanistically drive amyloidosis? In theory, they could either promote APP processing and Aβ generation or interfere with Aβ removal (or a combination of both). We first investigated APP metabolism in *Cnp*^*-/-*^ *5×FAD* mice. Axonal swellings that stain positive for APP are prominent features of ischemia, injury and myelin disorders and we speculated that these swellings contribute to the generation of Aβ in myelin mutant animals. Indeed, stalling axonal transport was shown to enhance amyloid production likely by increasing BACE1 and APP encounter in axonally transported vesicles^34–37^ and axons seem to be important sites of amyloid secretion^38–40^. Moreover, axonal swellings surrounding amyloid plaques (Fig.3a) are thought to be production sites of Aß^41,42^. Plaque-independent axonal swellings in *Cnp*^*-/-*^ mice are enriched in vesicular structures likely of endosomal/lysosomal origin (Fig.3b). Using antibodies against APP processing enzymes and multiple APP/Aβ-specific antibodies (Fig.3a), we found that axonal swellings in *Cnp*^*-/-*^ *5×FAD* brains accumulate β- and γ-secretase and consequently stain positive for β- and γ-cleaved APP fragments and Aβ (Fig.3c,d). We validated these findings by Western blot analysis of white matter and cortex and found elevated levels of BACE1 (statistically significant in white matter) (Fig.3e) in 6-month-old *Cnp*^*-/-*^ *5×FAD*. When we investigated APP processing by immunoblotting, cortical APP metabolism was shifted towards an increased abundance of CTF-β and CTF-α fragments, but without changes to full length APP abundancy or the α/β CTF ratio (Fig.3f). Together, these findings suggest that myelin defects enhance Aß generation.

**Figure 3.**
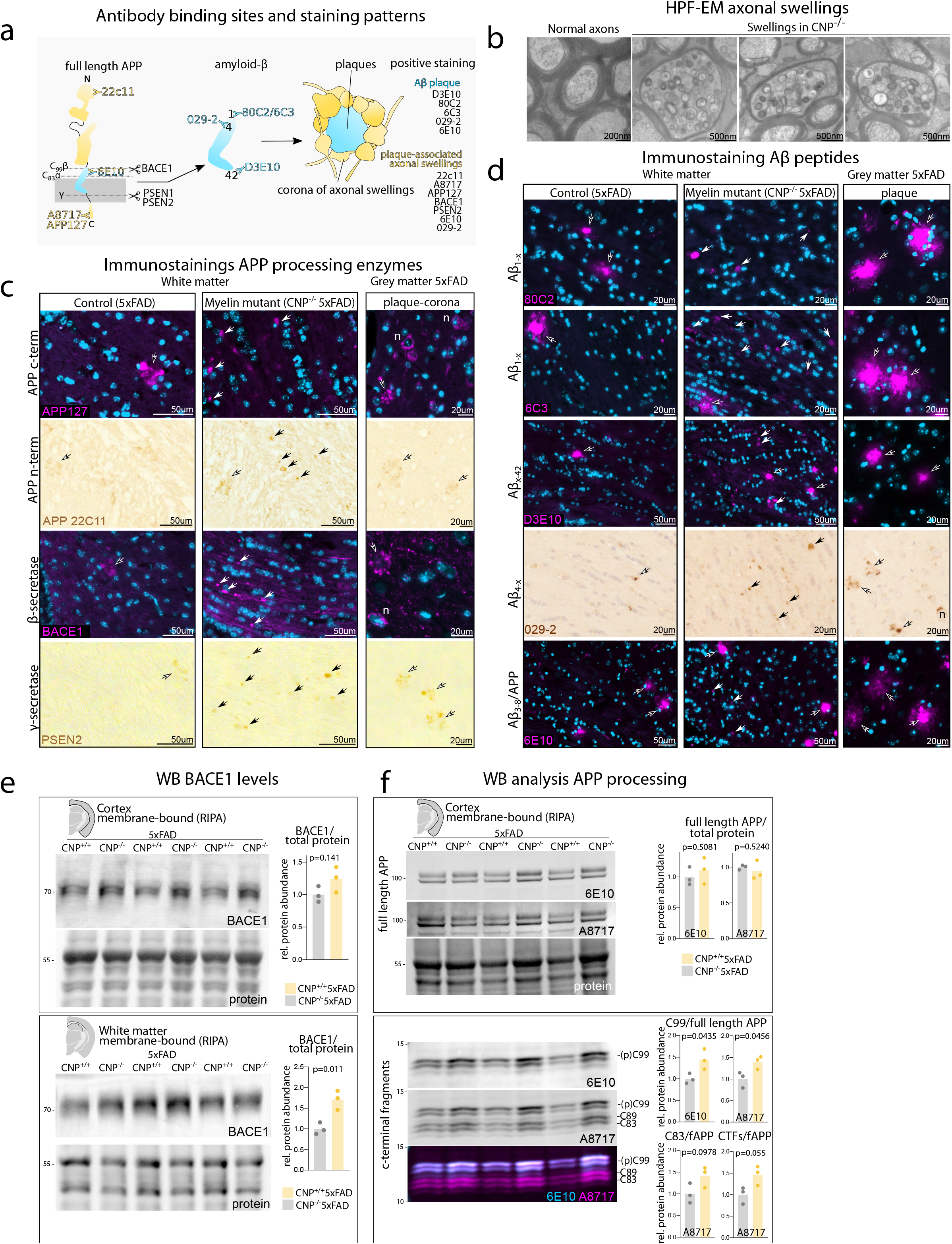
Myelin dysfunction alters APP processing. **(a)** Schematic representation of binding sites and specificity of antibodies to APP and amyloid-β used in the following panels and typical staining observed (plaque-associated axonal swellings vs Aβ plaque staining). Plaque-associated axonal swellings typically form a corona consisting of multiple swellings around amyloid plaques. **(b)** High pressure freezing EM of optic nerves from 6-month-old wildtype and CNP^-/-^ optic nerves. Examples of (plaque-independent) axonal swellings in CNP-deficient animals show abundant accumulation of endosomal/lysosomal structures and multivesicular bodies. **(c)** Fluorescent and chromogen immunostainings against APP and APP cleavage enzymes BACE1 (β-secretase) and Psen2 (as part of the γ-secretase complex) showing accumulation of APP, BACE1 and PSEN2 in plaque-independent axonal swellings in CNP^-/-^ 5×FAD. Contoured arrows mark plaque-associated axonal swellings typically forming a corona as abundantly found in the cortex of 5×FAD mice. Non-contoured arrows indicate plaque-independent axonal swellings as observed in CNP^-/-^mice. **(d)** Fluorescent and chromogenic immunostainings against different species of Aβ-peptides in CNP^-/-^ 5×FAD vs 5×FAD in the white matter. As additional control, typical plaque staining in 5×FAD cortex are shown. Contoured arrows indicate proper amyloid plaques, typically stained very intensely. Non-contoured arrows indicate swellings stained positive by the respective β-amyloid antibody, typically less intensely stained and of round structure. D3E10, 80C2 and 6C3 do not show cross-reactivity to full length APP and typically do not stain plaque-associated swellings, but stain swellings in CNP^-/-^ 5×FAD mice. 029–2 and 6E10 antibody show certain cross-reactivity to full-length APP and also stain axonal swellings. **(e)** Fluorescent immunoblot analysis of BACE1 levels in micro-dissected cortex and white matter of CNP^-/-^ 5×FAD and 5×FAD mice. P-value of unpaired, two-sided Student’s t-test is indicated in the respective quantification bar plots. Dots represent lanes/single animals. n=3 for each group. **(f)** Fluorescent immunoblot analysis of APP fragmentation in the membrane-bound fraction of micro-dissected cortical tissue of CNP^-/-^ 5×FAD and 5×FAD control mice. Quantification of band intensity is given on the right. fAPP levels were normalised against total protein fastgreen staining. CTF levels were normalised to fAPP levels. fAPP: full length APP. CTFs: c-terminal fragments. P-values of unpaired, two tailed Student’s t-test is indicated in the respective quantification bar plots. Dots represent lanes/single animals. n=3 for each group.

### Myelin defects alter the microglial phenotype in AD mouse models

Glial cells play important roles in the clearance of myelin debris and amyloid peptides and microglia form barriers around amyloid plaques^43^. Thus, we investigated how microglia react to amyloid plaques when additionally challenged with defective myelin. Despite pronounced gliosis throughout the brain in both *Cnp*^*-/-*^ *5×FAD* and *Cnp^-/-^APP*^*NLGF*^ mice, we noticed that cortical microglia failed to cluster around amyloid plaques (Fig.4a,b). Such a phenotype has been previously described in *Trem2* loss of function scenarios^43–45^. We therefore asked if myelin dysfunction interferes with the upregulation of *Trem2* and the induction of the disease-associated microglia (DAM) signature, that is associated with a plaque corralling phenotype^46,47^. Using magnetic activated cell sorting (MACS)-isolation of microglia and RNA sequencing (Fig.4c), principal component analysis (PCA) revealed that the transcriptome of *Cnp*^*-/-*^ *5×FAD* microglia was dominated by the *Cnp*^*-/-*^ signature (Fig.4d). Extraction of the top 100 genes contributing to PC1 revealed 4 major gene clusters with different trajectories throughout the experimental groups (Fig.4e, Extended data Fig.6a). Homeostasis markers (*Cx3cr1, P2ry12, Tmem119*) that were downregulated in 5×FAD microglia were even further downregulated in *Cnp*^*-/-*^ and *Cnp*^*-/-*^ *5×FAD* mice (Fig.4e,f). Conversely, DAM signature genes (*Clec7a, Gpnmb, Apoe, Spp1, Axl* and *Itgax*) were upregulated in *5×FAD* microglia, and even further increased in *Cnp*^*-/-*^ and *Cnp*^*-/-*^ *5×FAD* microglia (Fig.4e,f, Extended data Fig.6c) likely increased fraction of reactive microglia in the *Cnp*^*-/-*^ brain. Importantly, *Trem2* and *Tyrobp* showed an unaltered induction in *Cnp*^*-/-*^ *5×FAD* mice (Fig.4e,f). We noticed that *Apoe*, a well-known factor in Aβ aggregation^48^, was massively upregulated in *Cnp*^*-/-*^ and *Cnp*^*-/-*^ *5×FAD* which is likely to lead to elevated APOE protein levels in the brain parenchyma of these mice. Indeed, using immunostainings, we found elevated levels of APOE in white matter and amyloid plaques in *Cnp*^*-/-*^ *5×FAD* (Fig.4g).

**Figure 4.**
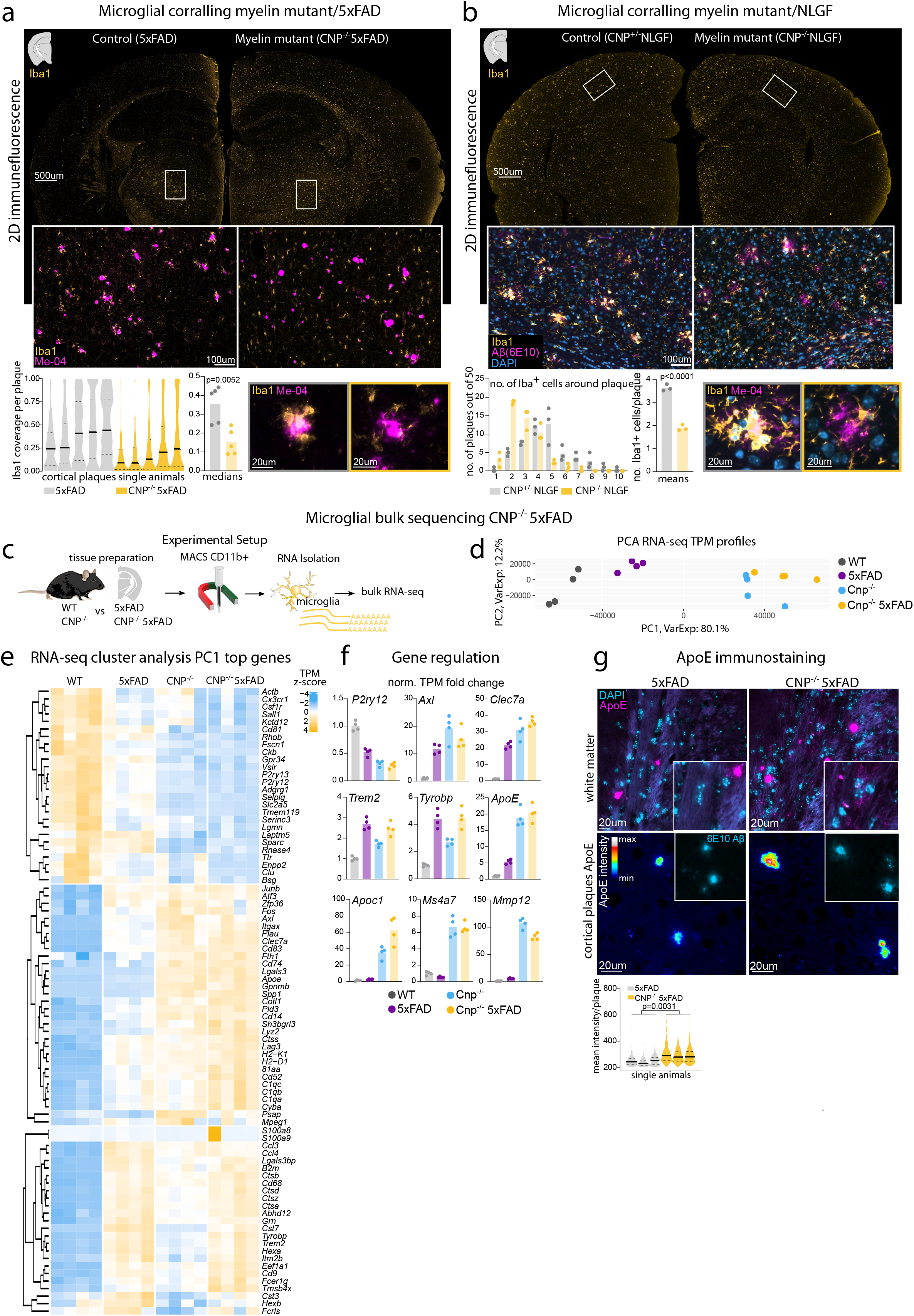
Loss of microglial corralling around amyloid plaques in myelin mutant mice. **(a)** 2D immunofluorescence analysis of microglial reaction to amyloid plaques in CNP-/-5×FAD and 5×FAD mice by Iba1 and amyloid co-staining. Automated quantification of Iba1 plaque coverage in the cortex. Each violin plot represents a single animal. 2017 individual cortical plaques were analysed in 5×FAD, 2190 in CNP^-/-^ 5×FAD brain slices. Bar plots represent the median of each individual animal shown in the violin plots. Unpaired, two-tailed Student’s t-test p-value is given in the bar plot. n=5 per group. **(b)** 2D immunofluorescence investigation of microglial reaction to amyloid plaques in CNP^-/-^ NLGF and NLGF control mice by Iba1 and amyloid co-staining. Histogram distribution of the number of microglia per amyloid plaque in 50 representative plaques in CNP^-/-^ NLGF and NLGF mice. Dots represent single animals and bar represents mean values. Unpaired, two-tailed Student’s t-test p-value is given in the bar plot. n=3 per group. **(c)** Experimental setup for microglia bulk RNA-seq. Microglia were isolated from hemispheres (without olfactory bulb and cerebellum) of 6-month-old wildtype (WT), CNP^-/-^, 5×FAD and CNP^-/-^ 5×FAD animals and subjected to RNA-seq (replicates n=4 for each genotype). **(d)** Principal component analysis (PCA) was used for evaluating relative distances between normalised RNA TPM profiles. Dots represent replicates. PC1 explained 80.1% of data variability, and strongly reflected CNP^-/-^ dominated microglia transcriptome changes. **(e)** Heatmap of top genes contributing to PC1 variability. **(f)** Bar plots of selected genes (homeostatic marker *P2ry12,* disease-associated microglia signature T*rem2, Tyrobp, Axl, Clec7a* and differential regulated genes in CNP^-/-^ 5×FAD (*Apoc1, ApoE, Ms4a7, Mmp12*). Bar show normalised expression level. **(g)** Microscopic analysis of APOE levels in CNP^-/-^ 5×FAD and 5×FAD by immunofluorescence staining against ApoE. Upper panel shows strong APOE staining in amyloid plaques on both 5×FAD and CNP^-/-^ 5×FAD brains and speckled staining throughout the white matter in CNP^-/-^ mice indicative of elevated ApoE levels. Lower panel shows representative images of APOE plaque content in cortical plaques in 5×FAD and CNP^-/-^5×FAD mice, pseudo-coloured according to rainbow lookup table. Quantification shows violin plots of mean ApoE plaque content per plaque. Each violin plots represents a single animal (n=3 per genotype, 703 plaques for 5×FAD, 846 plaques for CNP^-/-^ 5×FAD). For statistical analysis, unpaired, two-tailed Student’s t-test was performed.

The interpretation of bulk RNA-seq data is limited. To better understand the DAM-like microglial signature in *Cnp*^*-/-*^ mutants, we performed single nuclei RNA-sequencing (snRNA-seq) on cortical and callosal tissue from *Cnp*^*-/-*^ and wildtype mice (Fig.5a, Extended data Fig.7a,b). Cluster analysis identified 4 major microglia subpopulations, including one with high expression of DAM marker genes (*Trem2, Lpl* and *Spp1*), which was only seen in *Cnp*^*-/-*^ mice (Fig.5b,c,d, Extended data Fig.7d). We next assessed similarities between this *Cnp^-/-^*-associated DAM cluster and 5×FAD-associated DAM by integrating our dataset with the microglial snRNA-seq profile of Zhou et al^44^ (Fig.5e). Indeed, *Cnp*^*-/-*^ DAM clustered with 5×FAD DAM (Fig.5e). Average expression analysis at the single cell level revealed that most DAM marker genes (including *Apoe, Lyz2* and *Axl*) were expressed at a similar level (Fig.5f). However, some genes (including *Ms4a7, Gpnmb* and *Lpl*) were highly induced in *Cnp*^*-/-*^ DAM, but only moderately so in 5×FAD DAM (Fig.5f). Importantly, *Trem2* was robustly upregulated in 5×FAD and to a lesser extent in *Cnp*^*-/-*^ DAM, suggesting that Aβ pathology is a stronger inducer of TREM2/TYROBP signalling (Fig.5f). We noted that genes of the *Ms4a* family were highly induced in microglia of *Cnp*^*-/-*^ 5×FAD mice, which is also a feature of some microglia in early development^49^. The same *Ms4a* genes regulate the level of soluble TREM2 and modify AD risk^50^. However, immunoblot analysis ruled out that myelin dysfunction affects TREM2 protein simply via increased *Ms4a* expression (Extended data Fig.8).

**Figure 5.**
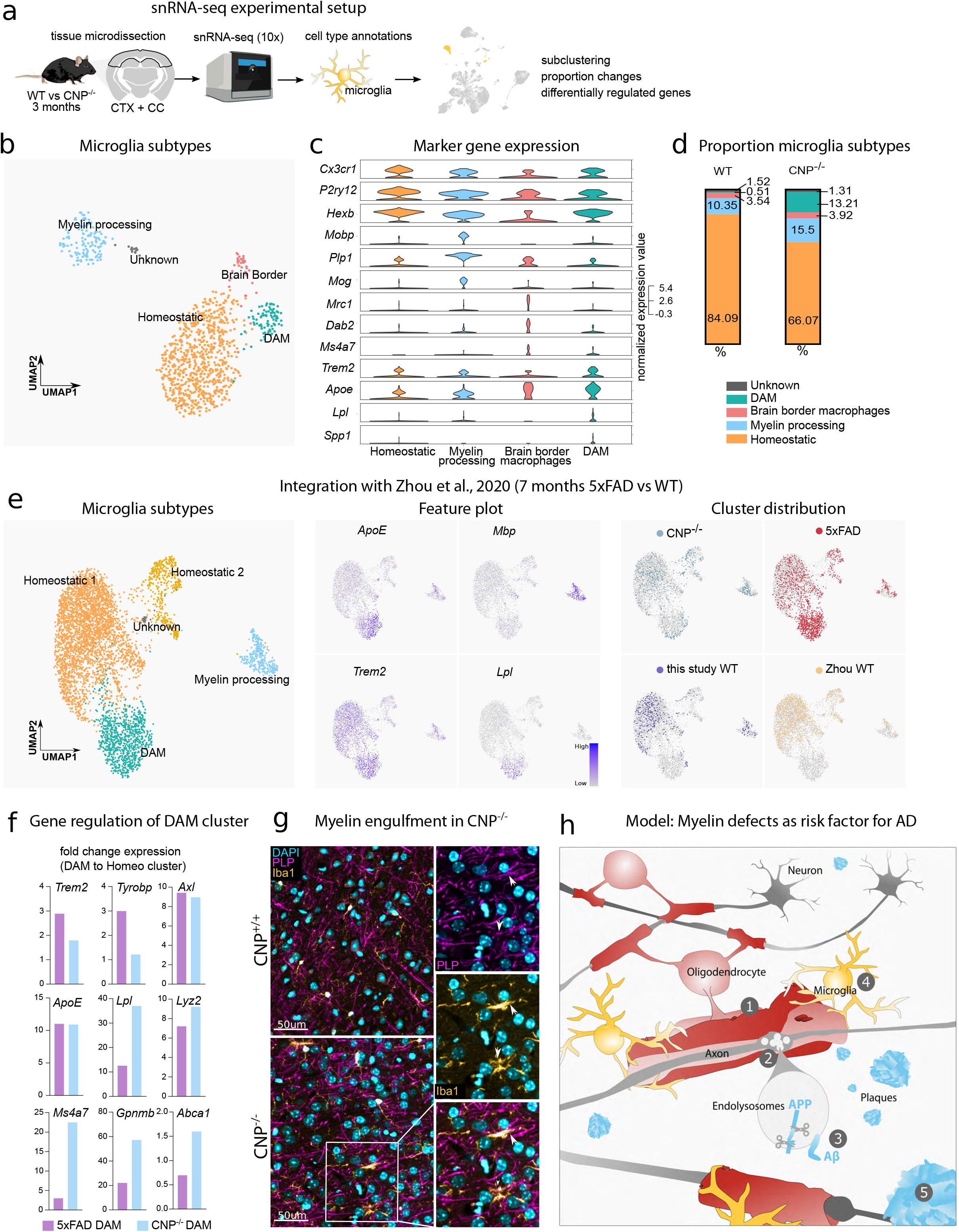
Myelin dysfunction induces DAM-like state as determined by snRNA-seq and distracts microglia from plaques. **(a)** Experimental setup for studying myelin-dysfunction induced microglia profiles at the single cell level. Cortex and adjacent callosal white matter were micro-dissected from 3-month-old wildtype and CNP^-/-^ mice and subjected to snRNA-seq (10x). Cell types were identified based on marker genes and microglia were subset for further analysis. **(b)** UMAP visualization of annotated microglia subtypes. **(c)** Violin plots showing marker gene expressions in each microglia subpopulation. **(d)** Bar plot shows distribution of microglia subpopulations in wildtype and CNP^-/-^. **(e)** Integrative analysis of this study and 7-month-old 5×FAD and wildtype microglia snRNA-seq profiles from Zhou et al.^38^ Feature plots showing normalised expression levels of ApoE, Mbp,Trem2 and Lpl for identification of microglia subclusters. Right panel showing genotype distribution among microglia subpopulations visualised in the UMAP space. **(f)** Bar plots representing average regulation levels of single genes in CNP^-/-^ DAM and 5×FAD DAM. Regulation values are calculated by comparing gene average expression of DAM to the respective homeostatic microglia population. **(g)** Immunofluorescence analysis of myelin phagocytosis (PLP+) by microglia (Iba1+) in the cortex of CNP^-/-^ mice. **(h)** Scheme illustrating a model of AD in which myelin dysfunction (1) induces axonal spheroids (2) where endo/lysosomes accumulate and Aβ production is enhanced (3). Simultaneously, microglia become increasingly engaged with myelin and are distracted from amyloid plaques (4). All processes contribute to enhanced formation of amyloid plaques (5).

In summary, our transcriptomic data suggest that microglia challenged by myelin dysfunction are still genetically equipped to react to amyloid plaques, but surprisingly fail to do so *in vivo*. We thus propose that microglia, once engaged in the clearance of defective myelin, are distracted from amyloid plaques. Indeed, *Cnp*^*-/-*^ microglia strongly express genes involved in phagocytosis (*C1q, C3, Itgax*), lysosomal function (*Spp1, Gpnmb, Lyz2, Pld3, Rab7b*) and lipid handling (*ApoE, Lrp, Lpl, Msr1, Apoc1, Apobec1, Abca1*) (Fig.4e,5f, Extended Data Fig.6c). Moreover, in Cnp mutants we validated by immunolabelling and the colocalization of PLP and IBA1 that microglia actively engulf myelin membranes (Fig.5g). Likewise, our snRNA-seq analysis identified a subpopulation of microglia that showed high level of “myelin RNA” (*Mobp, Plp1, Mbp, Mog* transcripts), which was further increased in *Cnp*^*-/-*^ microglia (Fig.5c,d, Extended data Fig 7c). This cell cluster likely represents microglia that had phagocytosed myelin (including oligodendroglial mRNA), as recently reported for snRNA-seq of multiple sclerosis autopsies^51^. We surmise that such microglial distraction not only leads to a faster build-up of amyloid in the brain, but that factors secreted by activated yet myelin-engaged microglia (such as APOE) may further fuel plaque seeding.

### Ageing myelin is a risk factor for AD

Based on our findings, we propose a resulting working model for AD, in which myelin of the ageing forebrain well known to lose structural integrity^2^ causes microglia to become engaged (as modelled in myelin mutant mice). This microglial activity interferes with the ability to clear Aβ deposits and prevent plaque formation (Fig.5h). Simultaneously, ageing myelin loses its axon-supportive functions, which increases neuronal BACE1 and APP-CTF levels suggestive of enhanced Aβ production (Fig.5h). It is tempting to speculate that other well-known AD risk factors such as traumatic brain injury or cardiovascular diseases convey risk via negatively impacting myelin health or at least through similar mechanism i.e. microglia ‘distraction’.

A critical role of myelin deterioration in the progression of AD has been previously hypothesised in light of correlative macroscopic brain imaging studies^52^, but necessarily lacked any evidence of causality. Our experimental findings in genetic models of AD can provide such a causal link with a molecular underpinning, and are fully in line with the prevailing amyloid hypothesis^9^ and the emerging role of neuroinflammation in disease progression^53^. Moreover, we can position the well-documented loss of myelin integrity in the ageing primate brain as the most upstream initiator of AD pathology, which helps to explain why age is the major AD risk factor. Our working model is further supported by classical neuropathological observations by Braak & Braak^54^ who pointed out the conspicuous temporal relationship with CNS myelination, which occurs the latest in those brain regions that are the first to develop AD pathology. Mechanistically, cortical ensheathment by oligodendrocytes is thin and may lack the robust myelinic channel architecture found in subcortical tracts that appear resistant to AD pathology and may overcome the axon shielding effects of myelin by providing glial metabolic support.

Only few studies have experimentally investigated oligodendrocytes and myelin in AD^55–60^. Nearly all have focussed on the effect of amyloid plaques and Aβ on oligodendrocytes and myelin, i.e., revealing secondary de- and hypomyelination. Similarly, scRNA-seq studies using human AD autopsy samples have identified myelin-related transcripts among the top altered clusters in AD patients^44,61^. These findings are likely the downstream effects of overt Aβ and tau pathology on oligodendrocytes and their precursor cells in the latest stages of disease. Our discovery that age-dependent loss of myelin integrity can be a driver or risk factor of amyloid deposition not only changes our view on the role of oligodendrocytes in AD, together with the downstream effects of Aβ pathology causing demyelination, this closes a “vicious cycle” in the AD brain that may explain why a long prodromal phase is followed by a more rapid clinical progression. If further corroborated in humans, promoting myelin health should be considered as a target for the delay or prevention of AD.

## Material and Methods

### Human tissue analysis

Case selection was performed from a pool of approximately 400 individuals, in which an autopsy with neuropathological evaluation was performed between 2018 and 2019 as a matter of routine procedure following death at the Leipzig University Hospital. In the individual contracts that govern medical treatment, all patients provided upon admission written consent to the scientific use of tissue removed and stored after any biopsy or during autopsy. Selection of patients was performed according to exclusion/inclusion criteria. Samples were anonymized and processed in a blinded manner. Selected patients were of mixed age and gender, between >60 years and <90 years of age, had a clinical history of dementia, and a NIA-AA score in neuropathological assessment between A2B3C2 and A3B2C2 (inclusion criteria) and did not suffer from another severe neurological disorder (exclusion criteria). In addition, control patients of the same age range without any clinical or neuropathological record of neurological disease were selected. No other criteria besides the described characteristics were applied. In total paraffin-embedded CNS tissue of 3 patients with moderate to pronounced AD neuropathological changes according to the NIA-AA (see above) and 3 control patients was histologically evaluated. In our analysis we used biopsies from the medial temporal lobe containing the hippocampal formation. We performed histological assessment for intracortical myelin content on the human tissue provided. For this, we sectioned paraffin blocks (5μm) and stained paraffin-sections for CNP, PLP, Iba1 and amyloid plaques simultaneously (see Histology for details).

### Mouse strains and husbandry

Animal experiments were conducted in compliance with german animal welfare practices and approved by the local authorities (Landesamt für Verbraucherschutz und Lebensmittelsicherheit, Niedersachsen). Mice were group-housed in the local animal facility of the Max Planck Institute for Experimental Medicine under a 12-h dark/12-h light cycle and fed *ad-libitum*. Both sexes were used throughout the study. Original mouse strains used to generate crossbreeedings were: 5×FAD^62^, APP^NLGF 63^, CNP^-/- 23^, PLP^-/y 24^, MBP^fl/fl 64^, Emx-Cre^65^. For analysis, either littermate controls were used or a corresponding control line from the initial F1 generation was generated. Age of animals is given in the respective figure legend. All animals were maintained on a C57BL/6 background. Genotyping was performed on clips derived from earmarking according to standard protocols (see references for original mouse strains). Genotype was confirmed by regenotyping on a tail biopsy upon sacrifice of the animal at the end of the respective experiment.

### Demyelination models

As demyelination models, we employed cuprizone-intoxication and experimental autoimmune encephalomyelitis (EAE) and experiments were conducted as described previously^66^. For cuprizone-mediated demyelination, male 14 weeks old 5×FAD mice were fed a powder diet containing 0.2% (w/w) cuprizone (Sigma-Aldrich) for 4 weeks followed by a 4-weeks recovery period on a normal pelleted food without cuprizone supplementation. Control 5×FAD received a standard diet without cuprizone. Animals were perfused at 18 weeks of age and brain tissue was analysed via light sheet microscopy and epifluorescence microscopy. For EAE experiments, 10-week-old 5×FAD mice were immunised against myelin oligodendrocyte glycoprotein (MOG) by subcutaneous injection of 200mg MOG peptide 35–55 in complete Freund’s adjuvant (Mycobacterium tuberculosis at 3.75 mg/ml; BD) followed by injection of 500ng of pertussis toxin (Sigma) at day 1 and 3 post EAE induction (dpi 1 and 3). Animals were checked on a daily basis and a neurological disease score was determined according to the following parameters: : 0 - normal; 0.5 - loss of tail tip tone; 1 - loss of tail tone; 1.5 - ataxia and mild walking deficits (slip off the grid); 2 - mild hind limb weakness, severe gait ataxia and twist of the tail causing rotation of the whole body; 2.5 - moderate hind limb weakness and inability to grip the grid with the hind paw but ability to stay on an upright tilted grid; 3 - mild paraparesis and falls from an upright tilted grid; 3.5 - paraparesis of the hind limbs (legs strongly affected but move clearly); 4 - paralysis of the hind limbs and weakness in the forelimbs; 4.5 - forelimbs paralyzed; 5 - moribund or dead. Animals were perfused at 28 dpi at 14 weeks of age. All immunised 5×FAD animals developed EAE (Extended data Fig.3d). Brain and spinal cords were analysed by light sheet microscopy and epifluorescence microscopy.

### Mouse behavioural testing

Hindlimb clasping was assessed by suspending the mouse on its tail for 5sec, carefully observing movement of the hindlimbs and scoring movement impairments according to a score from 0–4: 0 – no impairments, hindlimbs normally spread and moved; 1 - one hindlimb shows slightly less mobility; 2 - both hindlimb show less mobility, 3- both hindlimbs show reduced mobility and reduced spread; 4- both hindlimbs show severely reduced movement and severely reduced spread. Videos were recorded. Animals were tested in the elevated plus (EPM) and Y-maze (YM) on consecutive days. In the EPM paradigm, animals were allowed to freely explore an elevated cross-shaped platform with two opposing enclosed arms and two opposing open arms for 5min without the experimentator present. In the YM paradigm, animals were allowed to explore a Y-shaped maze with enclosed arms for 8min without human interference. Videos were recorded and analysed using the Bioobserve Viewer behavioural analysis setup with automated tracking. Zones (Open arms, closed arms, centre) and object detection settings were optimised according to the maze type used. After the run, positional data, track length and full track images were exported. For the EPM, the time spent in both open arms was summed and plotted. For the YM, the order of arm entries was recorded, and the number of correct triads (consecutive visit of the three different arms) determined. The alternation index was calculated according to the formular alternation index = number triads/number of arm entries-2. For a successful arm visit, animals had to enter an arm with their full body (excluding tail).

For statistical analysis of the behavioural data, we report the results of several different type III ANOVAs that probed the main effects for the 5×FAD and myelin mutants as well as their interaction. All analyses were conducted in R (version 4.04, R Core Team 2021). The analyses of variance were computed with the afex package as reported in ^67^.

### Tissue preparation

For microscopic analysis, animals were deeply anesthetised or euthanised via CO2 asphyxiation and subsequently transcardially perfused with ice-cold Hank’s buffered salt solution (HBSS) and 4% paraformaldehyde (PFA) in 0.1M phosphate buffer, pH 7.4. Brains and spinal cord were removed and postfixated overnight in 4% PFA/phosphate buffer. Brains were washed once in PBS pH7.4 and stored in PBS at 4°C until further use. For biochemical analysis of full brains, animals were killed by cervical dislocation, brain and spinal cord were extracted and fresh-frozen on dry-ice. Tissue was stored at −80°C until further use. For microdissection of subcortical white matter and cortical tissue, animals were sacrifised by cervical dislocation and the brain was quickly removed and submerged in ice-cold HBSS. The brain was inserted into a custom-made brain matrix and the brain was sliced coronally in ~1mm thick slices. Brain slices were spread onto an ice-cold glass plate and cortex and subcortical white matter were separated and excised from each individual brain slices. Tissue was immediately frozen on dry-ice and stored at −80°C until further use.

### Whole tissue staining and clearing for light sheet microscopy

Fixed hemibrains and spinal cords were pretreated and permeabilised following a modified iDisco protocol^26,27^. Briefly, tissues were dehydrated with an ascending concentration of methanol in PBS (50% 1x, 80% 1x, 100% 2x, 1 h each). Tissues were then bleached with a 1:1:4 ratio of H_2_O_2_:DMSO:methanol overnight at 4°C. Further dehydration followed with 100% methanol incubation in 4°C (30 min), −20°C (3 h), and overnight storage at 4°C. Samples were incubated the following day in methanol with 20% DMSO before subjecting them to gradual rehydration in a descending methanol in PBS series (80% 1x, 50% 1x, 0% 1x, 1 h each). We then washed the tissues with a detergent mixture of 0.2% Triton X-100 in PBS (2x, 1 h) and permeabilised them overnight at 37°C in PBS with 0.2% Triton X-100, 20% DMSO, and 0.3 M glycine. After permeabilization, tissues were either stained with the Congo Red dye (Sigma Aldrich) for beta sheet structures within amyloid plaques or immunolabelled with antibodies of interest. For Congo Red staining, tissues were washed with PBS/0.2% Tween-20/10 mg/ml heparin/5mM sodium azide (PTwH) (2x, 1 h) before immersing them for 3 d at 37°C in 0.005% w/v Congo Red (100x stock solution in 50% ethanol). For immunolabelling, following tissue permeabilization and glycine treatment, samples were blocked in PBS with 0.2% Triton X-100, 10% DMSO and 6% goat serum (GS) for 3 d followed by washing in PTwH (2x, 1 h) and incubation in primary antibodies with the appropriate dilution factors (1:500, rabbit, anti-Iba, Wako) in PTwH with 0.2% Triton X-100, 5% DMSO and 3% GS for 14 d at 37°C. Upon completion of primary antibody labelling, tissues were washed in PTwH (6x, 10 min) and stored overnight at 37°C. For secondary antibody labelling, tissues were again incubated in secondary antibodies with the appropriate dilution factors (anti-rabbit Dylight 633, 1:500, Thermo-Fisher) in PTwH with 3% GS for 7 d at 37°C. Prior to clearing, spinal cords were fixed in 1.5% w/v Phytagel in water. Dyed or immunolabelled tissues were washed in PTwH (3x, 10 min each) before rehydration through an ascending concentration of methanol in PBS (20% 1x, 40% 1x, 60% 1x, 80% 1x, 100% 1x, 1 h each) and delipidation in a 1:2 mixture of methanol:dichloromethane (1x, 1 h 40 min). Lastly, samples were cleared by immersing them in ethyl cinnamate (Eci, Sigma-Aldrich) until transparent. All incubation steps were carried out at constant medium-speed rotation at the indicated temperatures. Samples were stored at room temperature in Eci until imaging.

### *In toto* imaging of whole tissues and analysis/visualization

Cleared hemibrains and spinal cords were imaged *in toto* with a light sheet microscope setup (UltraMicroscope II, LaVision Biotec) equipped with a 2x objective lens, zoom body, and a corrected dipping cap. Samples were imaged being submerged in a sample chamber containing Eci. For all hemibrain imaging, hemibrains were mounted medial side-down on the sample holder to acquire sagittal images. Images were acquired in the mosaic acquisition mode with the following settings: 5 μm light sheet thickness, 20% sheet width, 0.154 sheet numerical aperture, 4 μm z-step size, 1000×1600 px field of view, 4×4 tiling, dual light sheet illumination, and 100 ms camera exposure time. Red fluorescence was recorded with 561 nm laser excitation at 80% laser power and a 585/40 emission filter. Far-red fluorescence was recorded with 640 nm laser excitation at 30% laser power and a 680/30 emission filter. Autofluorescence was recorded with 488nm laser excitation at 50% and 525/20nm emission filter. Images were imported into Vision4D v3.2 (Arivis) and stitched using the tile sorter setup. Partly, images were imported and stitched using the Imaris Importer and Stitcher v9.1 (Bitplane). Rendered whole hemibrains were then processed and three main regions of interest (ROIs) were manually annotated based on the sagittal Allen mouse brain atlas, namely isocortex, hippocampus, and alveus. All ROIs were first traced manually in 2D planes to automatically extrapolate the 3D ROIs. Cortical and hippocampal annotations were cropped with a medial cut-off of approximately ~0.4 mm and a lateral cut-off of ~4.4 mm which would span the dorsal isocortex and the entire hippocampal formation of one hemibrain. The lateral cut-off for the alveus ROI is the plane where the hippocampal formation appears in 2D. Next, we segmented amyloid plaques within the ROIs. For 3-month-old 5×FAD brain data, we typically used automated intensity thresholding. For 6-month-old 5×FAD data, plaques were segmented using the blob finder algorithm in Vision4D with the following parameters: 20 μm object size, 10–15% probability threshold, and 0% split sensitivity. Once segmentation has been performed, a stringent noise removal was performed by deleting objects with voxel sizes <10 from the object table. Objects were then critically reviewed and any additional noise, which might include but are not restricted to blood vessels and nuclei, were manually removed from the dataset. For quantification of amyloid plaques in the APP^NLGF^ brains that typically stained much weaker for Congo Red, plaque segmentation was performed using the machine learning segmenter in Vision4D. Object parameters, namely volume, ROI volumes, were extracted for further quantification. For plaque visualization in the different ROIs, objects are represented in centroids and color-coded according to location.

### Paraffin slices and histological stainings

Fixated hemibrains and spinal cords were subjected to dehydration steps (50% EtOH, 80% EtOH, 100% EtOH, 100% Isopropanol, 50% Isopropanol/50% Xylol, 2x 100% Xylol) followed by paraffinization on the STP 120 tissue processing machine (Leica microsystems). Samples were retrieved and embedded in paraffin blocks on the HistoStar embedding workstation (Epredia). Paraffin-embedded blocks were sectioned coronally while spinal cords were sectioned longitudinally at 5 μm slice thickness, slices were mounted onto slides and dried overnight. Slides were deparaffinised at 60°C followed by incubation in xylol (100% 2x) and a 1:1 mixture of xylol and isopropanol (1x) for 10 min each. The slides were rehydrated in a descending ethanol series. This was followed by incubation in either acidic antigen retrieval solution (pH 6.0) or basic antigen retrieval solution (10mM Tris/1mM EDTA pH10) (for 5 min and boiling for 10 min. The samples were cooled for 20 min and washed in distilled water for 1 min before a subsequent permeabilisation in 0.1% Triton X-100 in PBS. For Aβ antibody staining, samples were subjected to an extra antigen retrieval in 88% formic acid to loosen up the β-sheet structure for optimal antibody binding for 3min. For plaque ApoE staining, formic acid treatment was even prolonged (10min, fresh formic acid). This was followed by washes in PBS (2x, 5 min) and blocking with 10% GS in PBS for 1 h at RT. For chromogenic labelling, an additional step of inactivation of endogenous peroxidases was implemented prior to blocking by incubation in 3% H2O2. After blocking, slices were incubated in primary antibody solution (PBS/10%GS or PBS/5%BSA) overnight at 4°C in coverplates (Epredia). Antibodies used in this study were: anti-Iba1 (rabbit, Wako; 1:1000); anti-Aβ-6E10 (mouse, Biolegend; 1:1000), anti-CNP (mouse, AMAb91072, Atlas; 1:1000), anti-PLP-clone aa3 (rat, culture supernatant; 1:200), anti-BACE1 (rabbit, ab183612, Abcam; 1:100), anti-MBP (rabbit, serum, custom Nave Lab; 1:1000), anti-GFAP (mouse, GA5, Leica, 1:200), anti-n-terminal APP (22c11, Merck; 1:1000), anti-c-terminal APP (rabbit, 127–003, Synaptic Systems; 1:1000), anti-Aβx-42-D3E10 (rabbit, Cell Signalling Technology; 1:1000), anti-Psen2 (rabbit, Abcam, 1:100), anti-ApoE D7I9N (rabbit, Cell Signalling Technology; 1:500), anti-Aβ_1-x_ (mouse, 80C2, Synaptic Signalling, 1:200), anti-Aβ_1-x_ (mouse, Moab2-6C3^68^, Abcam, 1:200), anti-Aβ_4-x_ (guinea pig, 029–2^69^, Oliver Wirths, custom, 1:200). For immunofluorescence staining, samples were washed in PBS or Tris/2% Milk and incubated with the corresponding fluorescent secondary antibody diluted in PBS containing 10% goat serum for 2 h at RT in the dark. Fluorescently conjugated secondary antibodies used were: anti-mouse Alexa555 (donkey/goat, Thermo-Fisher; 1:1000), anti-mouse Dylight633 (donkey/goat, Thermo-Fisher; 1:1000), anti-rabbit Alexa555 (donkey/goat, Thermo-Fisher; 1:1000), anti-rabbit Dylight633 (donkey/goat, Thermo-Fisher; 1:1000). Amyloid plaques were stained by the β-sheet dye Methoxy-X04 (4 μg/ml in 50% ethanol, Tocris) for 30 min at RT and contrasting in 50% ethanol. Nuclei were either stained with DAPI (300nM, Thermo-Fisher) or ToPro3 (1:1000, Thermo-Fisher) in PBS for 5min at RT. Slides were again washed in PBS and mounted with Aqua PolyMount mounting medium (PolySciences). For chromogenic labelling, the LSAB2 Kit (Dako) for rabbit/mouse and the DAB-Kit from Zytomed was used according to the manufacturers’ protocols. Slides were then rinsed in water, dehydrated and mounted using Eukitt (Sigma-Aldrich).

### Epifluorescence and brightfield microscopy

Epifluorescence microscopy was performed on a Zeiss Observer Z1 microscope equipped with Plan-Apochromat 20x/08 and Fluar 2.5x/0.12 objectives, a Colibri 5 LED light source (630nm, 555nm, 475nm 385nm excitation), 96 HE BFP, 90 HE DAPI/GFP/Cy3/Cy5, 38 GFP, 43 DsRed, 50 Cy5 Zeiss filter sets, a Axiocam MrM and a SMC900 motorized stage. For whole brain slice microscopy, a preview scan at 2.5x magnification was taken and focus support points were distributed and manually set for imaging at 20x magnification in the ZEN imaging software (Zeiss). Tiled images were stitched in ZEN. For visualisation, pseudocolours (cyan, magenta, yellow) were assigned to different channels, intensity was adjusted and images were exported in ZEN. Brightfield microscopy of DAB-stained slices was performed on a Zeiss Axiophot Imager.Z1 equipped with a Achroplan 4x/0.1, PlanFluar 20x/0.75 and Plan Neofluar 40x/0.75 objects and a AxioCamMrc camera. For whole brain slice microscopy, a preview scan at 4x magnification was taken and focus support points were distributed and manually set for imaging at 40x magnification in the ZEN imaging software (Zeiss). Tiled images were stitched in ZEN. Brightness and contrast of RGB images were adjusted and images exported in ZEN.

### 2D microscopy quantification

2D image analysis was performed in FIJI (v1.53c)^70^. For quantification of amyloid load, thresholding was performed to segment amyloid-β deposits (either stained by Methoxy-04 or amyloid-β plaques) and area positive was calculated in the respective region of interest. For analysis of the plaque corralling phenotype, in Iba1+/Aβ co-stainings individual plaques were segmented using thresholding and the Iba1+ area in each individual plaque was calculated using a FIJI macro-script. Iba1 coverage was expressed as percentage. For quantification of ApoE levels in plaques, ApoE positive plaques were segmented and the raw mean fluorescence per plaque was calculated.

### High-pressure freezing electron microscopy

Sample preparation by high pressure-freezing and freeze substitution was performed as described ^71^. In brief, optic nerves of 6-months-old CNP^-/-^ and control wildtype mice were freshly dissected, immersed in 20% PVP in PBS and placed into HPF sample carriers (Wohlwend, Sennwald, Switzerland). After freezing using a HPM100 high pressure freezer (Leica Microsystems, Vienna, Austria), samples were embedded in EPON after freeze substitution using 0.1 % tannic acid in acetone followed by 2% OsO4 and 0.1 % uranyl acetate in acetone. After polymerization, samples were sectioned with a UC7 ultramicrotome (Leica Microsystems, Vienna, Austria) and imaged with a LEO912 transmission electron microscope (Carl Zeiss Microscopy GmbH, Oberkochen, Germany) using an on-axis 2k CCD camera (TRS, Moorenweis, Germany).

### Cell fractioning

For analysis of APP and Trem2 fragmentation, cell fractioning was performed prior to western blot analysis according to published protocols^45^. In brief, tissue was homogenised in DEA-buffer (0.25% Diethylamine 50mM NaCl pH10) using the Precellys bead-milling method (Precellys soft tissue homogenising lysis kit, Bertin Instruments) and the soluble protein fraction was extracted via centrifugation (10min, 500g, 4°C) followed by ultracentrifugation (1h, 130 000g, 4°C). The membrane-bound fraction was solubilised in RIPA buffer (20mM Tris-HCl pH7.5, 150mM NaCl, 1%NP-40, 1%SDS, 2.5mM sodium pyrophosphate, 1mM Na_2_EDTA) and cleaned via centrifugation (10min, 500g, 4°C) and ultra-centrifugation (1h, 130 000g, 4°C). RIPA insoluble material (containing plaque Aβ) was resuspended in ice-cold 70% formic acid in water, sonicated and ultracentrifuged (1h, 130 000g, 4°C). Supernatant was collected as formic acid fraction and neutralised with 1M Tris pH9.5. All buffers and solutions were supplemented with protease inhibitor cocktail (P8340, Merck). Fractions were stored at −80°C until further use.

### SDS PAGE and Western blotting

To determine sample protein concentration, detergent compatible protein assays (Biorad) were carried out in duplicates. Sample was mixed with Laemmli sample buffer (2%SDS, 10% glycerol, 0.0025% bromphenol blue, 0.125M Tris-Cl, pH 6.8, 0.05M DTT) and equal amount of protein (typically 20–30ug) were loaded per lane. For BACE1 western blotting, standard Tris-Glycine SDS-PAGE gels (8%) were used. For western blot analysis of APP and TREM2 fragmentation, Tris-Tricine SDS PAGE gels (10-20%, Novex, Thermo Fisher) were used. Gels were run at 100–120V for approximately 1h. For Tris-Glycine SDS-PAGE gels, proteins were transferred onto low fluorescent Immobilon-FL membrane (0.45μm por size, Merck) using the Biorad wet-blot system (1.15h, 500mA) and blot transfer buffer (25mM Tris, 190mM glycine) containing 20% MeOH. For Tris-Tricine SDS PAGE gels, proteins were transferred onto a low fluorescent PVDF membrane of lower pore size (0.2μm). Blots were washed in water and transferred protein was stained with Fastgreen for transfer quality-check and for normalisation purposes. For this, membranes were transferred to Fastgreen working solution (0.0005% Fastgreen FCF (Serva) in de-staining solution (30% Methanol, 7% mL glacial acetic acid, 63% H2O) for 5min and briefly washed two times in de-staining solution. Membranes were imaged at a ChemoStar fluorescent imager (Intas) equipped with a 670 nm/20 nm excitation filter and near infrared emission collection. Membranes were rinsed in TBS with Tween (0.05%) until pH was neutral and blocked in 5% BSA in TBS-T for 1h at room temperature. Membranes were then incubated in primary antibodies in 5% BSA over night at 4°C on a rotating shaker. The following primary antibodies were used: anti-BACE1(1:1000, rabbit, D10E5, Cell Signalling Technologies), anti-n-terminal APP (1:1000, mouse, 22c11, Merck), anti-c-terminal APP (1:1000, rabbit, A8717, Merck), anti-APP/Aβ (1:1000, mouse, 6E10, Biolegend), anti-c-terminal TREM2 (1:1000, rabbit, E7P8J, Cell Signalling Technologies). Membranes were washed several times and membranes were incubated in secondary antibody solution (5% BSA in TBS-T). The following secondary antibodies were used: anti-rabbit IgG (H+L) DyLight 800 (1:1000, Thermo Fisher), anti-mouse IgG (H+L) Dylight 680 (1:1000, Thermo Fisher). For visualisation, membranes were scanned at an Odyssey platform (Licor). For quantification, background was subtracted from raw images and bands were analysed using FIJI (integrated density). Protein levels were normalised to fastgreen whole protein or in the case of APP and Trem2 cleavage to full length protein.

### Magnetic activated cell sorting of microglia sorting and bulk RNA-sequencing

Microglia were isolated from mouse hemibrains (excluding cerebellum and olfactory bulb) via magnetic-activated cell sorting. Dissected tissues were enzymatically and mechanically dissociated using the Miltenyi Biotec adult brain dissociation kit according to manufacturer’s protocol. Prior to microglial isolation via CD11b microglia microbeads and LS columns (Miltenyi Biotec), astrocytes (ACSA-2 microbeads) and oligodendrocytes (O4 microbeads) were removed to enhance purity of the microglial population. Isolated microglia were directly eluted in RLT lysis buffer and RNA was isolated using the RNeasy Micro Kit (Qiagen). In total, n = 4 replicates were used for each genotype (WT, Cnp^-/-^, 5×FAD, Cnp^-/-^ 5×FAD. RNA extracted from sorted mouse brain hemisphere microglia was eluted in 30 μl nuclease-free water and subjected to 50 bp single-end mRNA sequencing using HiSeq 4000 (Illumina). Raw sequencing data were first evaluated by FASTQC v0.72 for quality, then aligned against the reference mouse genome GRCm38 using STAR v2.5.2b-2^72^ with default parameters. Gene raw counts of each sample were extracted using featureCounts v1.6.3^73^ from aligned profiles for differential gene expression (DGE) analysis using DESeq2 v.1.26.0^74^ and converted to TPM value for sample distance calculation and visualization, as well as for gene expression pattern analysis. For DGE analysis, each pair of genotypes were calculated separately, and statistics results were summarised (SuppTable2). Gene targets with adjusted P-value < 0.05 were considered as significantly regulated. Using normalised TPM profiles, samples were embedded by principal component analysis (PCA) to assess distances.

### Nucleus isolation and single-nuclei transcriptome sequencing

Cortex and corpus callosum from 3-month-old Cnp^-/-^ and WT controls were micro-dissected and subjected to single-nuclei transcriptome sequencing. For each genotype and replicate, the tissue of two animals were pooled. Two replicates per genotype were sequenced. Nuclei were isolated according to previously published methods^75^. Briefly, frozen tissue was transferred into 2ml of pre-chilled homogenization buffer (320mM sucrose, 0.1% NP40, 0.1mM EDTA, 5mM CaCl_2_, 3mM Mg(Ac)_2_, 10mM pH 7.8 Tris, 167uM β-mercaptoethanol and 1x protease inhibitor (Roche)). Tissue was carefully homogenised and filtered through a 80μm strainer, and further centrifuged for 1min under 100 rcf. For each sample, 400 ul supernatant was collected into a pre-chilled 2ml low-binding Eppendorf tube, followed by adding 400ul 50% iodixanol solution (in 1x homogenization buffer containing 480mM sucrose) to reach a 25% iodixanol concentration. By layering 600ul of 29% iodixanol underneath the 25% iodixanol mixture, then 600ul of 35% iodixanol underneath the 29% iodixanol, two clear interfaces between different concentrations of buffers were created, and the tube was centrifuged for 20 min under 3000 rcf. After centrifugation, nuclei were collected from the band between the 29% and the 35% iodixanol layers and transferred to a fresh pre-chilled tube. Isolated nuclei were washed and resuspended in cold resuspension buffer (1xPBA, 1% BSA, 0.1U/ul RNase inhibitor) and further subjected to single-nuclei transcriptome libraries using the chromium single cell 3’ reagent kit according to the manufacturer's instruction (10x Genomics). The constructed libraries were sequenced using Novaseq 6000 (Illumina). Raw snRNA-seq data were collected in Fastq format and first aligned to the reference profile pre-mRNA (ENSEMBL GRCh38) using CellRanger toolkit v3.0.2 (10x Genomics).

Matrices containing UMI count of each gene in each nuclei were extracted for all samples, by filtering out nuclei with <200 detected genes and <500 total transcripts, as well as nuclei with outlier level transcripts quantity or gene detection rate identified according to individual sample sequencing depth (SuppTable3). Genes expressed in less than 3 cells were excluded for further analysis. Filtered expression matrices were combined, and UMI of each nucleus were normalised towards its total UMI counts with a scale factor of 10,000 and then log transformed.

### Dimensionality reduction, clustering analysis and cell type annotation

The normalised UMI matrix of all samples was mainly analysed using the R package Seurat v3.2.3^76^. High variable genes were calculated and scaled to support linear dimensionality reduction using PCA. For all cells, the first 50 PCs were used for further neighbouring embedding using uniform manifold approximation and projection (UMAP)^77^, as well as for performing the clustering analysis (resolution=0.5) using K nearest neighbour (KNN) algorithm. Cluster marker genes were calculated using the MAST algorithm^78^ to determine cluster cell type annotations. Clusters with undefined cell identities were removed from further analysis. To perform cell type specific analysis for microglia the corresponding cell population was firstly subset and reduced for its dimensionality using PCA. Similarly, selected top PCs were used for UMAP embedding and clustering analysis, with cluster marker genes calculated by MAST. Specific parameters used for analysing microglia can be found in (SuppTable3).

### External data integrative analysis

Integrative analysis was carried out between CNP^-/-^ and GSE140511^44^. More specifically, microglia from Cnp^-/-^ and corresponding controls were integrated with microglia profiles from 7-month-old WT, 5×FAD from GSE140511, to unravel the disease associated microglia (DAM) subpopulation. Integrative analysis was conducted using the SCTransform pipeline implemented in the Seurat toolkit. The batch effect corrected data underwent PCA dimensionality reduction, neighbouring embedding and unbiased clustering. Corresponding parameters for different datasets integration are listed in (SuppTable3).

### Data visualisation

Images were exported from the respective imaging or bioinformatic software (ZEN 2011 blue edition, Zeiss; Vision4D Arivis; FIJI; R) and final figures were assembled in Inkscape (v1.1, www.inkscape.org). Graphs were created in Prism 8.0 (Graphpad).

### Data availability

All raw sequencing data, as well as raw and processed counts matrices have been uploaded to the Gene expression Omnibus (GEO)^79^ under the SuperSeries accession number GSE178304. External datasets recruited for analysis were accessed through accession numbers listed in material and method.

### Code availability

Packages involved in data analysis pipelines are listed in material and method. The code used for bulk and single-nuclei transcriptome sequencing is available on GitHub https://github.com/TSun-tech/AD_MyelinMutant.git. More detailed information is available upon contacting the corresponding authors. Custom-made FIJI scripts for analysis of plaque-corralling are available on request.

## Acknowledgements

We thank the animal care takers and veterinarians of the animal facility at the Max-Planck-Institute for Experimental Medicine (MPI-EM). We thank Anette Fahrenholz and Katharina Overhoff for technical assistance. We thank members of the Department for Neurogenetics at the MPI-EM and the KAGS subgroup for helpful discussions and input. C.D. was supported by a Boehringer Ingelheim Fonds PhD fellowship. Work in KAN’s laboratory was supported by by the Dr. Myriam and Sheldon Adelson Medical Foundation (AMRF) and an ERC Advanced Grant (MyeliNANO).

## Author contributions

CD and KAN conceptualised and designed the study. CD and AOS performed microscopy and biochemical experiments. TSun planned and analysed bulk RNA-seq and snRNA-seq experiments. LS, SB, GS helped perform acute demyelination experiments. AAS helped analysing snRNA-seq experiments. SS provided mice. WM performed electron microscopic analysis and generated mice. SZ and OW helped analyse Aβ generation in axonal swellings. MT performed statistical analysis of behavioural data. TSaito and TSaido provided APP NLGF mice. DKB helped performing behavioural experiments. RK and DG performed snRNA-seq. MW and CH helped perform biochemical analysis of APP processing and provided critical experimental advice. RS provided human autopsy material. HE provided conceptual input. CD and TSun created figures. CD, TSun and KAN wrote the manuscript.

---

**Extended Data Figure 1.**
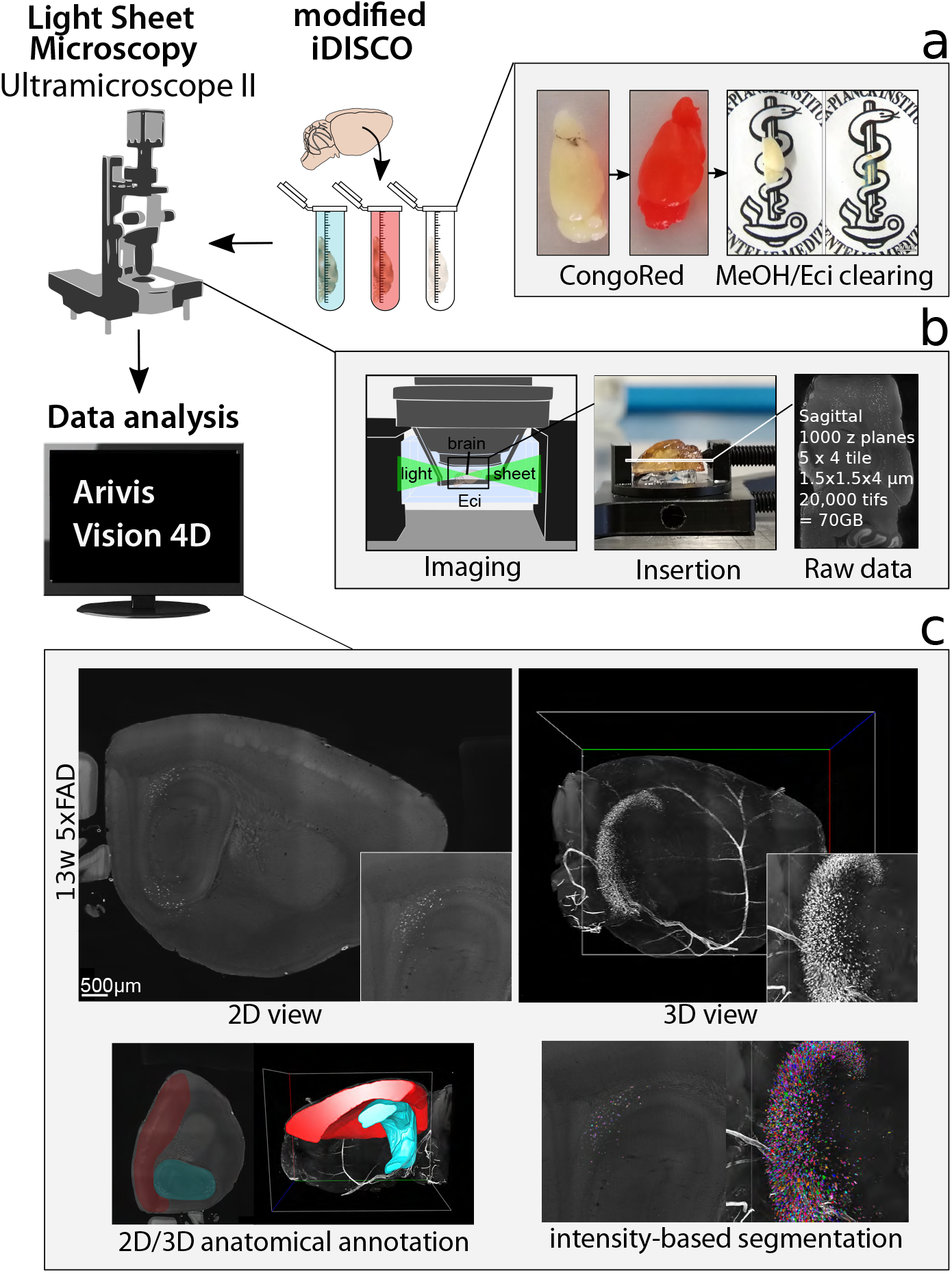
Assessment of in toto amyloid burden by light sheet microscopy. **(a)** Brains were subjected to *in toto* staining with the β-sheet dye Congo Red according to a modified iDisco protocol (see Material and Methods) followed by clearing in Ethylcinnamate (Eci). **(b)** Cleared brains were imaged on an Ultramicroscope II (LaVision-Biotech) light sheet setup to obtain sagittal optical slices. **(c)** Raw data were visualised and analysed in Arivis Vision 4D using manual region of interest annotation for hippocampus and cortex and automated plaque segmentation (intensity-thresholding: 3-month-old 5×FAD, blobfinder algorithm: 6-month-old 5×FAD, machine learning: 6-month-old APP^NLGF^).

**Extended Data Figure 2.**
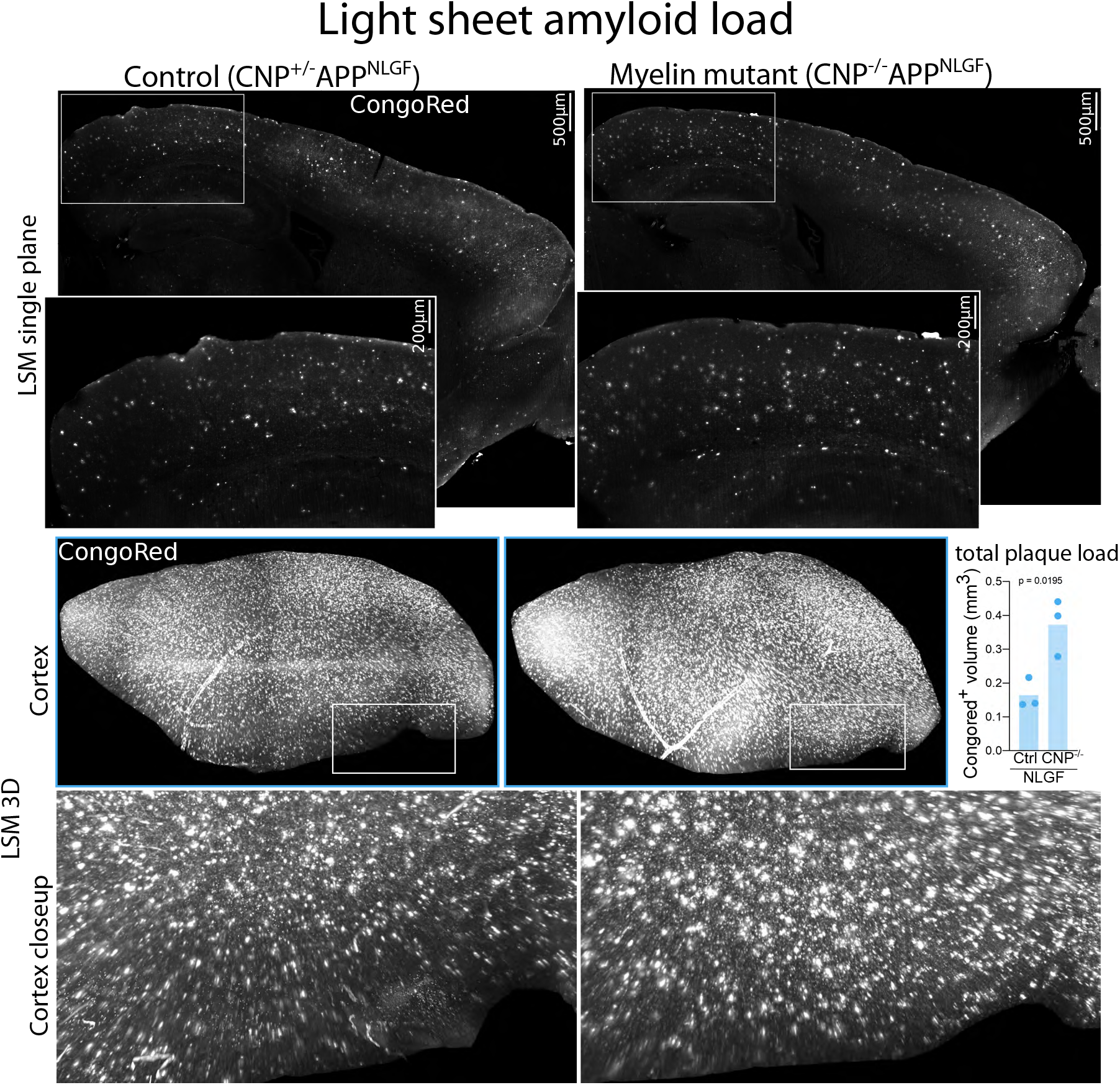
Light sheet microscopic analysis of amyloid plaque load in CNP^-/-^ APP^NLGF^. CNP^-/-^ APP^NLGF^ brains show enhanced plaque deposition at 6-month of age when compared to CNP+/-APPNLGF controls. Upper panel shows LSM single plane and a closeup of a cortical region. Middle panel shows 3D maximum intensity projection of the cropped isocortical region of interest. Plaque burden was quantified using machine-learning based segmentation of amyloid plaques. Lower panel shows the closeup indicated in the middle panel. Dots represent single animals and bars depict mean. Unpaired, two-tailed Student’s t-test was performed for statistical analysis.

**Extended Data Figure 3.**
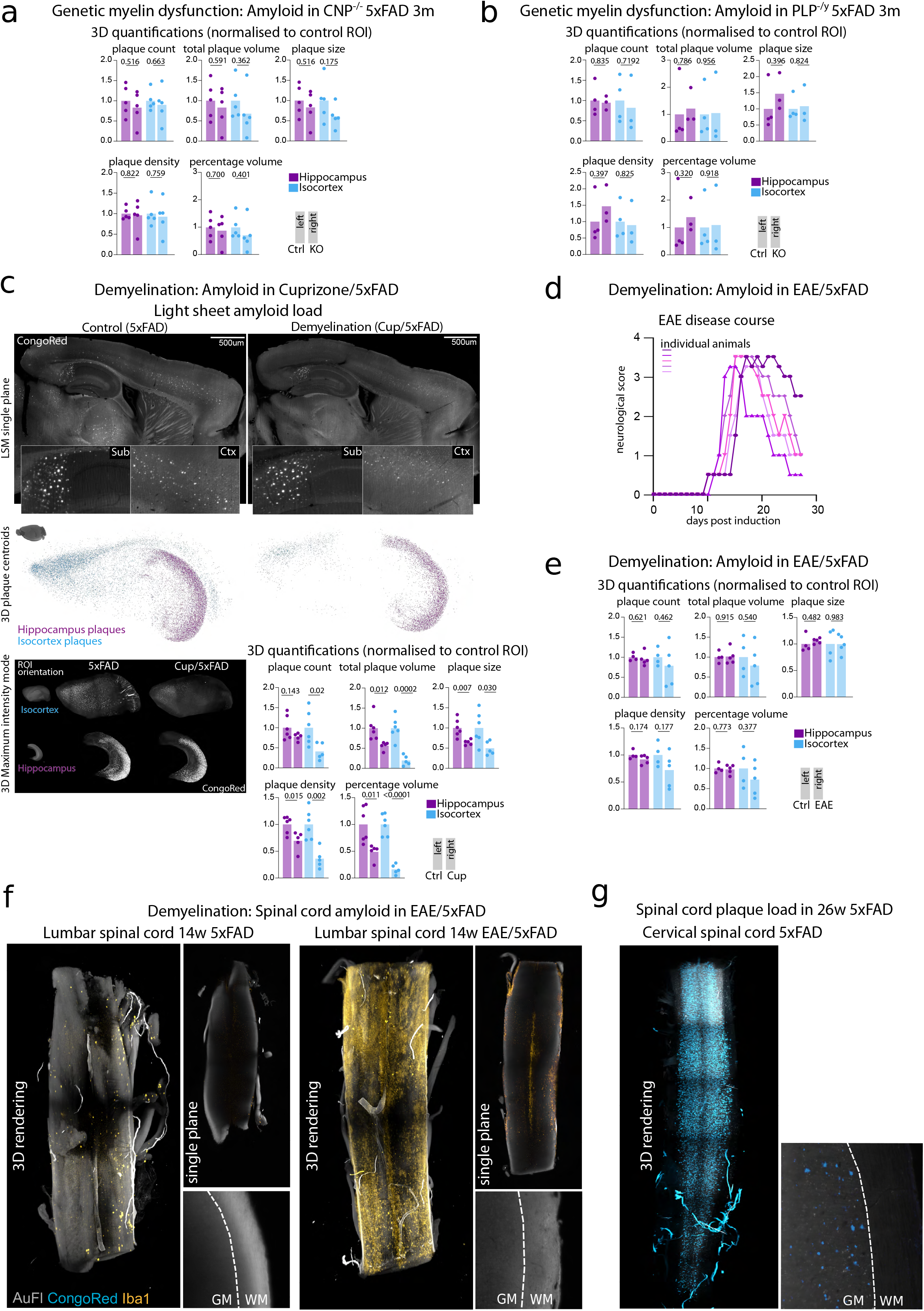
Microscopic analysis of plaque changes induced by dys- or acute demyelination (Main Figure 2 extended) **(a)** Quantifications of light sheet microscopic analysis of 3-months-old CNP^-/-^ 5×FAD mice. n=5 for control, n=5 for KO. **(b)** Quantifications of light sheet microscopic analysis of 3-months-old PLP-/y 5×FAD mice. n=4 for control, n=3 for KO. **(c)** Light sheet microscopic analysis of amyloid plaque load in cuprizone-fed 5×FAD mice. Upper panel shows LSM single plane of Congo Red stained brains and closeups of the subiculum and a cortical region. Middle panel shows 3D distribution of isocortical and hippocampal plaques represented as centroids. Lower panel shows 2D maximum intensity projection of cropped regions of interested. Lower right shows 3D quantifications of plaque burden parameters in hippocampus and isocortex. n=6 for control, n=5 for Cup. **(d)** Neurological scoring shows successful EAE-induction in 5×FAD animals with typical disease onset at around day 10 post induction. Lines represent single animals. n=5. **(e)** 3D quantifications of plaque burden in the brain of EAE/5×FAD animals. n=4 for control, n=5 for EAE. **(f)** Light sheet microscopic analysis of plaque burden in the lumbar spinal cord of EAE/5×FAD mice. No typical grey matter plaques could be detected in either 14 weeks old 5×FAD controls or EAE/5×FAD mice. The lumbar spinal cord was heavily affected by EAE lesions as visualised by Iba1 staining. **(g)** As positive control for successful detection of spinal cord plaques, a cervical spinal cord of a 6-month-old 5×FAD animal is shown. Throughout the panels, dots represent single animals and bars depict mean. Unpaired, two-tailed Student’s t-test was performed for statistical analysis.

**Extended Data Figure 4.**
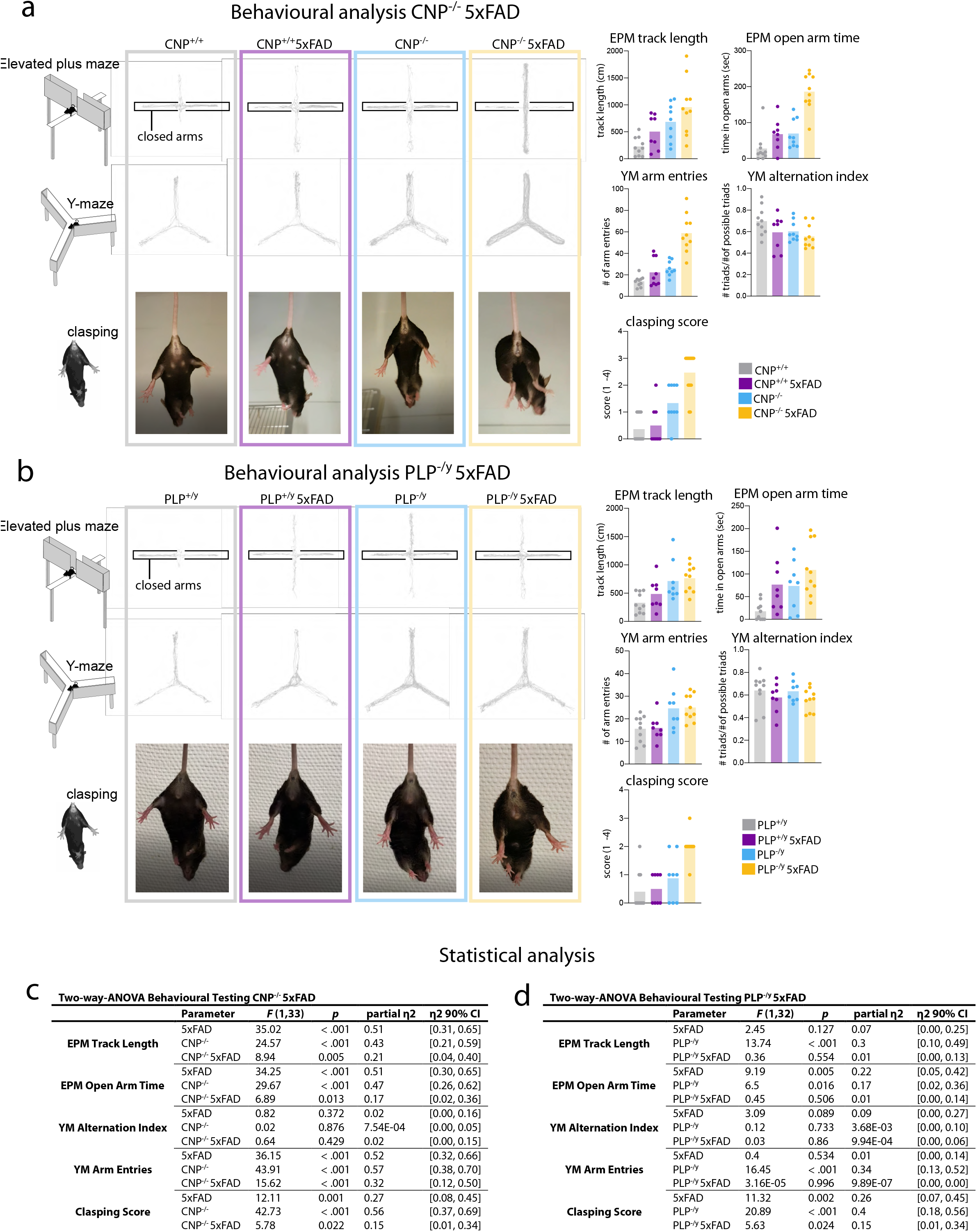
Myelin-dysfunction exacerbates behavioural deficits in 5×FAD mice. Behavioural analysis of mice in the elevated plus maze (EPM), Y maze (YM) and the clasping test. Middle panels show representative tracks in the respective maze and representative image of the clasping test. Right panel shows quantifications of behavioural parameters. Dots represent single animals, bars depict mean. **(a)** Behavioural analysis of CNP^-/-^ 5×FAD female mice. n=9 for CNP^+/+^, n=8 CNP^+/+^ 5×FAD, n=9 for CNP^-/-^, n=9 CNP^-/-^ 5×FAD **(b)** Behavioural analysis of PLP^-/y^ 5×FAD male mice. n=10 for PLP^+/y^, n=8 PLP^+/+^ 5×FAD, n=8 for PLP^-/y^, n=10 PLP^-/y^ 5×FAD **(c,d)** For statistical analysis, several different type III ANOVAs were performed for each behavioural test that probed the main effects for the 5×FAD genotype and the myelin-mutant genotype as well as their interaction.

**Extended Data Figure 5.**
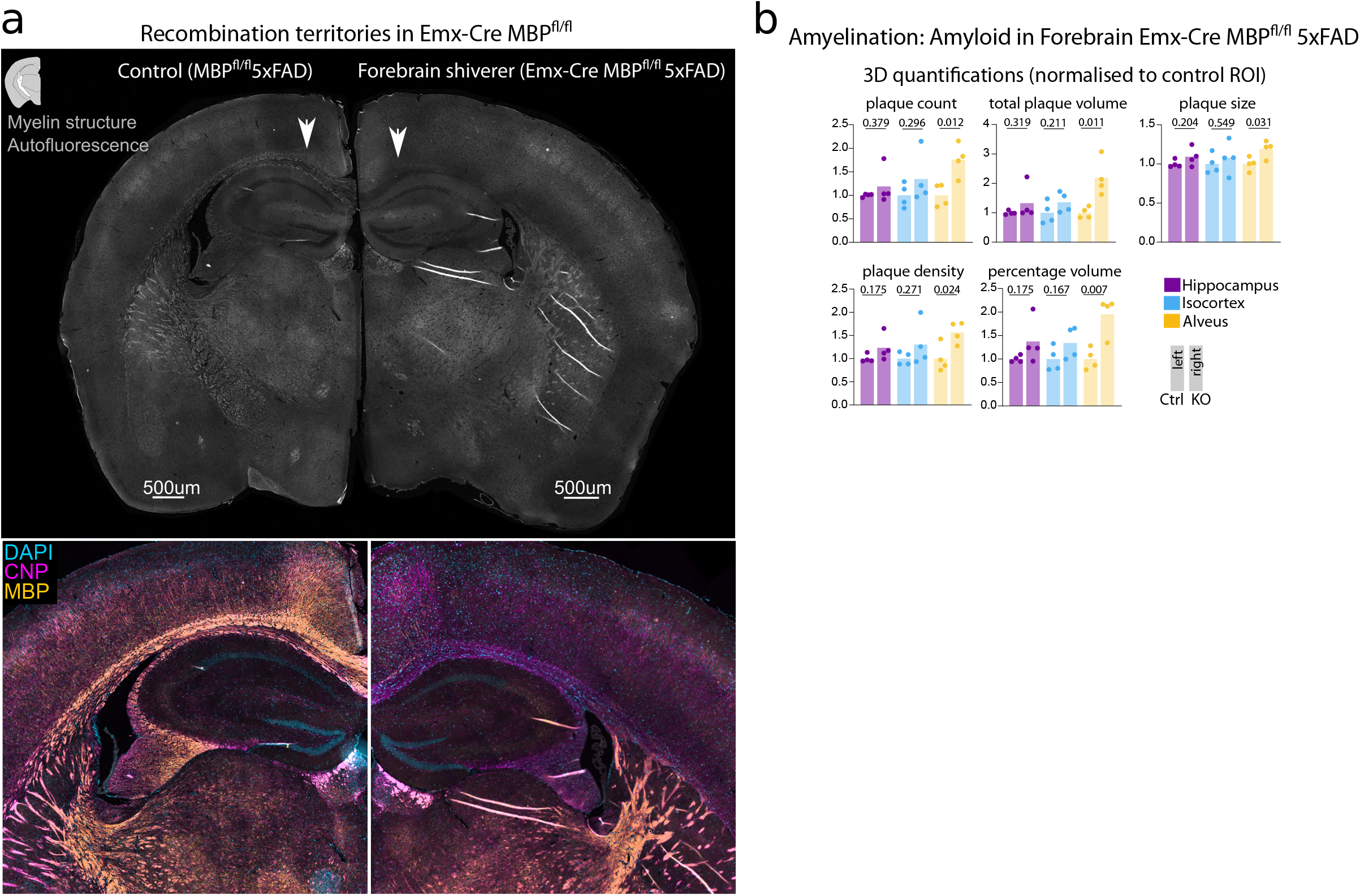
Recombination territories and plaque load in 6-month-old forebrain shiverer/5×FAD mice. **(a)** Basic characterisation of myelination patterns in Emx-Cre MBP^fl/fl^ 5×FAD mice. Autofluorescence shows clear lack of myelinated profiles in corpus callosum (arrows) while thalamus and striatum show normal myelin profiles. Lower panel show closeup images of anti-CNP and MBP co-immunolabelling in Emx-Cre MBP^fl/fl^ 5×FAD. Lack of myelin compaction (MBP-CNP^+^) throughout cortex, hippocampus and subcortical white matter. Compaction of myelin is unaffected in other brain regions such as thalamus and striatum (MBP^+^CNP^+^). **(b)** Quantification of light sheet microscopic analysis of plaque load in 6-month-old Emx-Cre MBP^fl/fl^ 5×FAD mice. Dots represent single animals, bar depicts mean. Umpaired, two-sided Student’s t-test was performed for statistical analysis for each ROI. n=4 for WT and cKO.

**Extended Data Figure 6.**
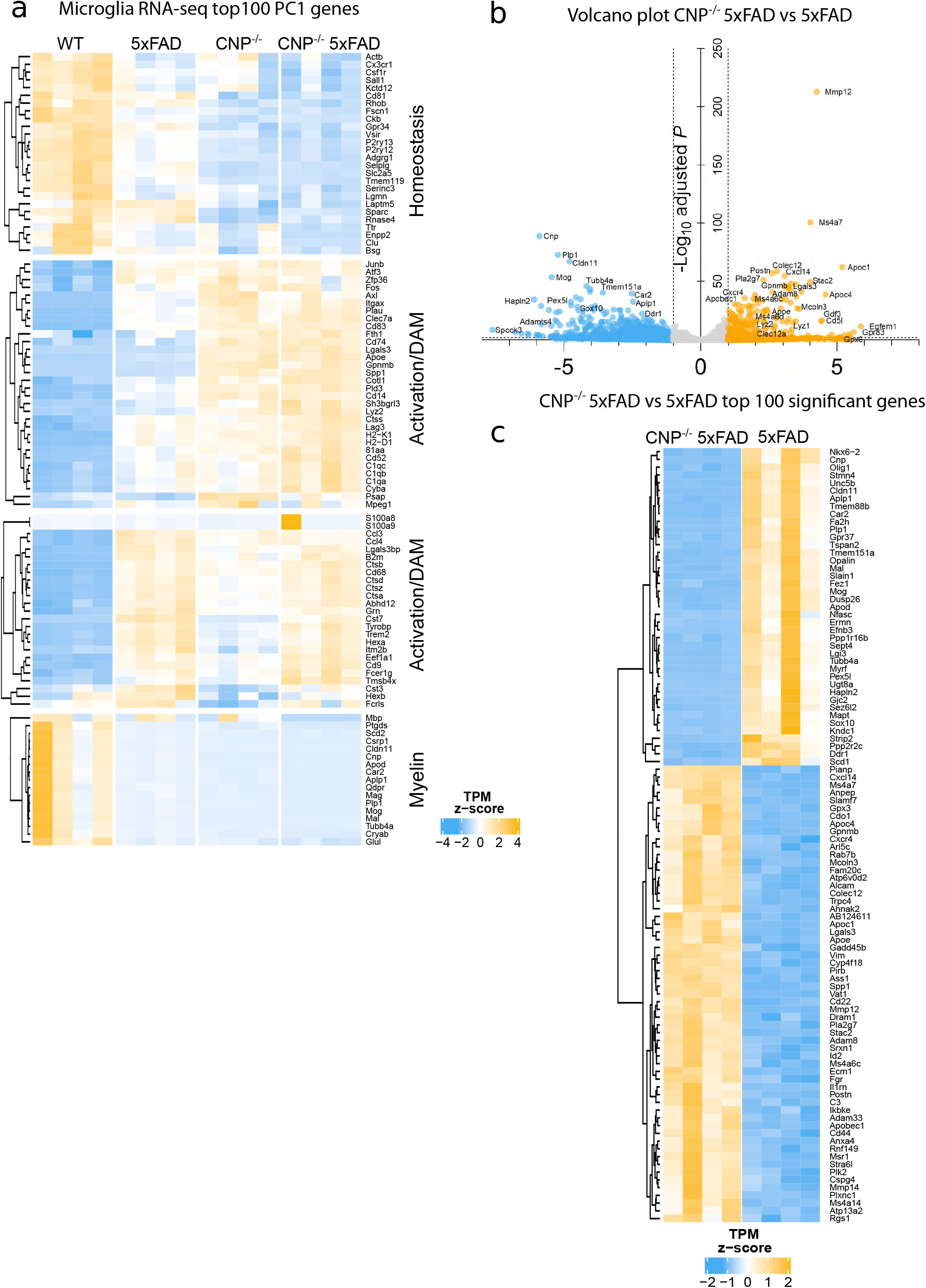
Transcriptomic signature of CNP-/- 5×FAD microglia (Figure 4 extended) **(a)** Heatmap of the top100 genes contributing to PC1 (Heatmap in Figure 4 extended). Four major clusters with different regulation trajectories were detected. **(b)** Volcano plot visualization of differentially expressed genes between CNP^-/-^ and CNP^-/-^ 5×FAD microglia (significant cutoff adjP < 0.01, log2FC > 1.0). **(c)** Heatmap represents normalised expression levels of top 100 significantly regulated genes between CNP^-/-^ 5×FAD vs 5×FAD microglia profiles as analysed by RNA-seq of isolated microglia.

**Extended Data Figure 7.**
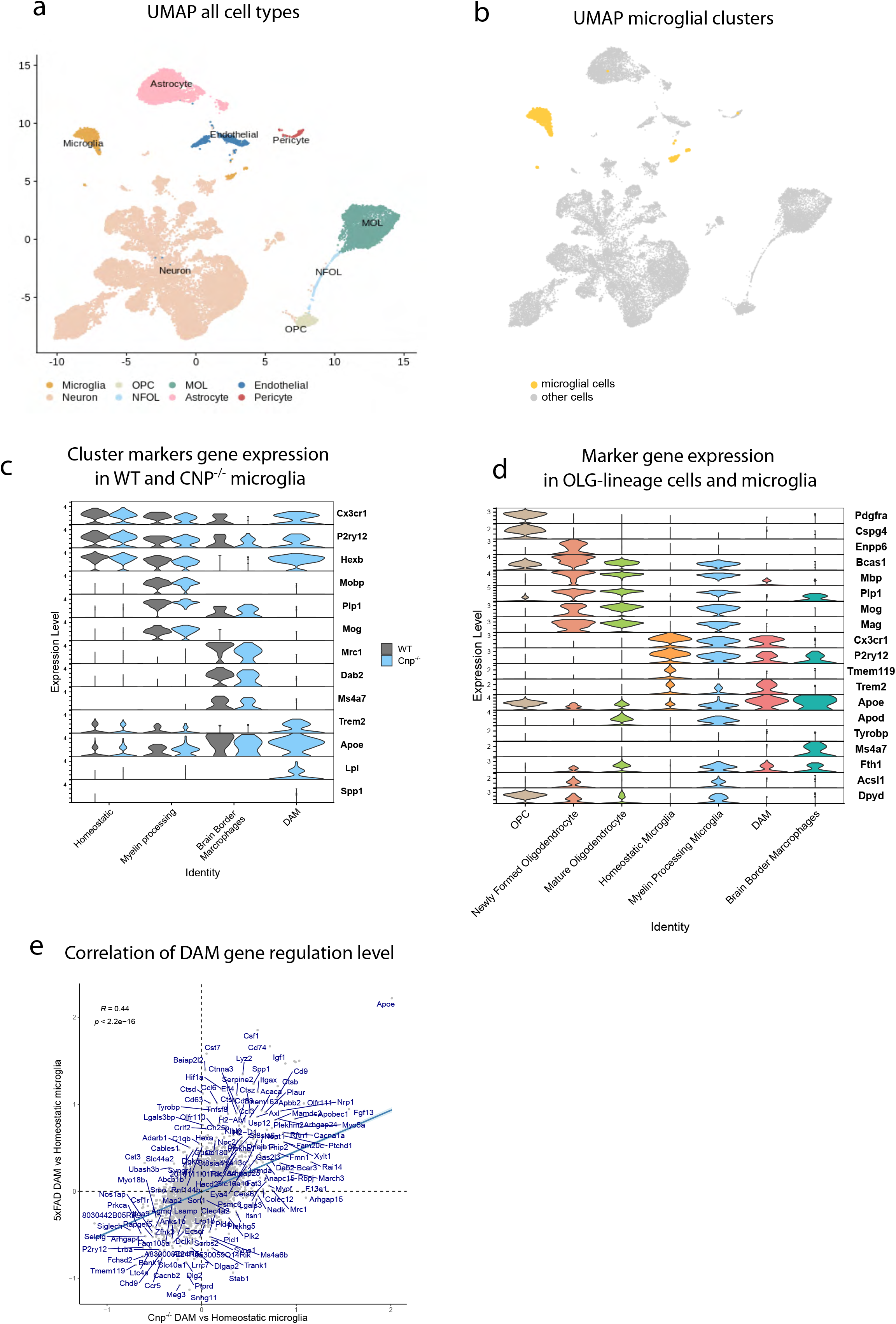
Single nucleus transcriptomic signature of CNP-/- brain (Figure 5 extended) **(a)** Celltype annotation in the UMAP space. **(b)** Microglial cells highlighted in the UMAP space (yellow). **(c)** Expression of marker genes for each microglial subcluster in CNP^-/-^ and WT mice. **(d)** Graph shows expression levels of oligodendroglial lineage cells and microglia marker genes throughout cell identities. Myelin processing microglia show robust presence of oligodendrocyte marker genes as well as microglial markers. **(e)** Scatter plot shows correlations of gene regulation levels between Cnp^-/-^ and 5×FAD DAM. Regulation levels were calculated by comparing gene average expression of DAM to homeostatic microglia in each genotype.

**Extended Data Figure 8.**
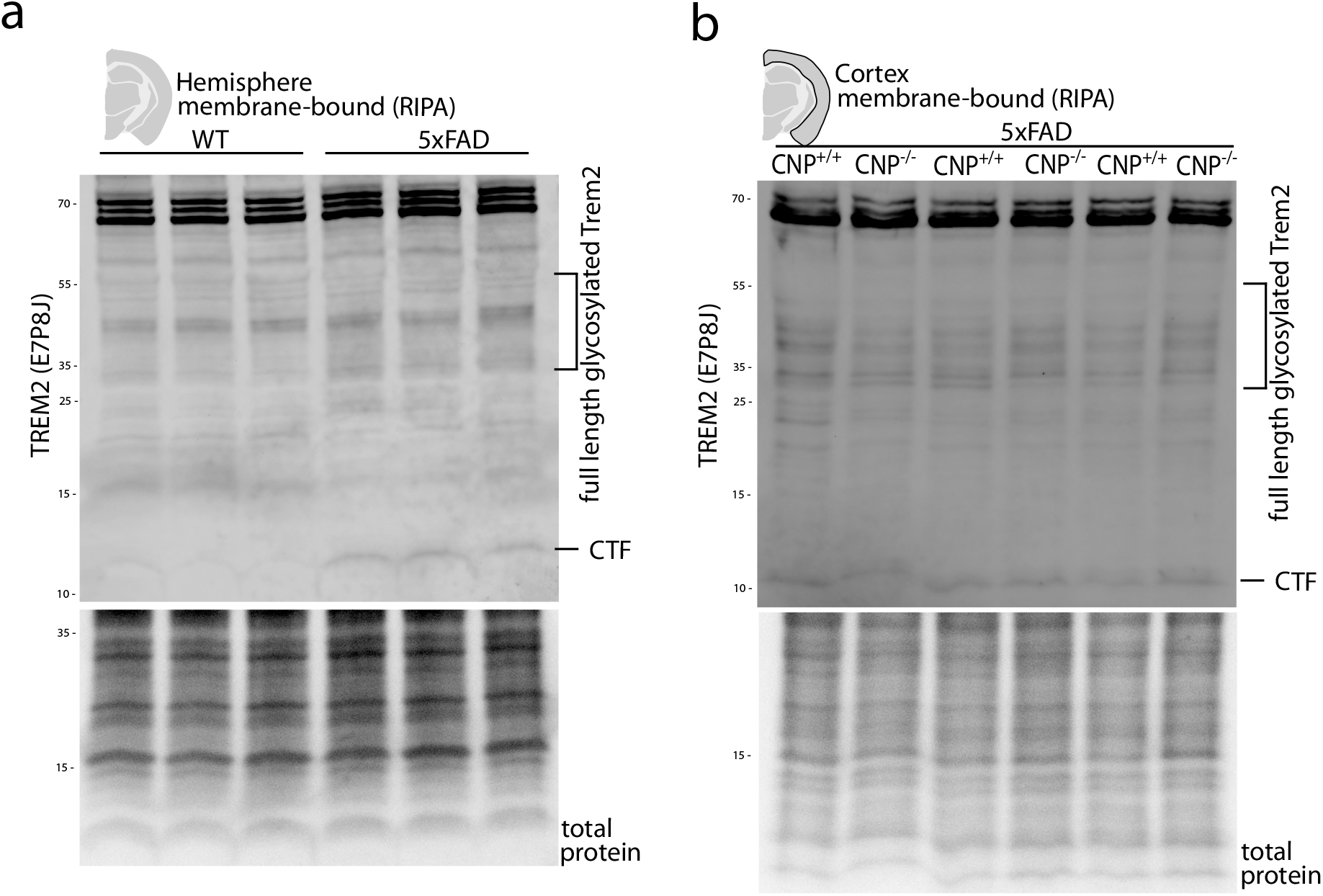
Trem2 cleavage is not altered in CNP^-/-^ 5×FAD mice. **(a)** Western blot analysis of Trem2 levels in WT vs 5×FAD mice shows clear induction of full length Trem2 (30-60kDa, glycosylated forms) and enhanced Trem2 cleavage (~12kDa) in 5×FAD mice. n=3 per group. Lanes present single animals. **(b)** Western blot analysis of Trem2 levels in CNP^-/-^ 5×FAD and 5×FAD mice shows no differences in Trem2 cleavage. Total protein fastgreen staining is shown as loading control. n=3 per group. Lanes present single animals.

## References

1 Shen, S., Liu, A., Li, J., Wolubah, C. & Casaccia-Bonnefil, P. Epigenetic memory loss in aging oligodendrocytes in the corpus callosum. Neurobiology of aging 29, 452–463 (2008).

2 Bowley, M. P., Cabral, H., Rosene, D. L. & Peters, A. Age changes in myelinated nerve fibers of the cingulate bundle and corpus callosum in the rhesus monkey. Journal of Comparative Neurology 518, 3046–3064 (2010).

3 Safaiyan, S. et al. Age-related myelin degradation burdens the clearance function of microglia during aging. Nature neuroscience 19, 995–998 (2016).

4 Griffiths, I. et al. Axonal swellings and degeneration in mice lacking the major proteolipid of myelin. Science 280, 1610–1613 (1998).

5 Fünfschilling, U. et al. Glycolytic oligodendrocytes maintain myelin and long-term axonal integrity. Nature 485, 517–521 (2012).

6 Lee, Y. et al. Oligodendroglia metabolically support axons and contribute to neurodegeneration. Nature 487, 443–448 (2012).

7 Mukherjee, C. et al. Oligodendrocytes provide antioxidant defense function for neurons by secreting ferritin heavy chain. Cell Metabolism 32, 259–272. e210 (2020).

8 Braak, H. & Braak, E. Neuropathological stageing of Alzheimer-related changes. Acta neuropathologica 82, 239–259 (1991).

9 Hardy, J. A. & Higgins, G. A. Alzheimer’s disease: the amyloid cascade hypothesis. Science 256, 184–186 (1992).

10 Hou, Y. et al. Ageing as a risk factor for neurodegenerative disease. Nature Reviews Neurology 15, 565–581 (2019).

11 Cohen, C. C. et al. Saltatory conduction along myelinated axons involves a periaxonal nanocircuit. Cell 180, 311–322. e315 (2020).

12 Saab, A. S. et al. Oligodendroglial NMDA receptors regulate glucose import and axonal energy metabolism. Neuron 91, 119–132 (2016).

13 Toyama, B. H. et al. Identification of long-lived proteins reveals exceptional stability of essential cellular structures. Cell 154, 971–982 (2013).

14 Ando, S., Tanaka, Y., Toyoda, Y. & Kon, K. Turnover of myelin lipids in aging brain. Neurochemical research 28, 5–13 (2003).

15 Lüders, K. A. et al. Maintenance of high proteolipid protein level in adult central nervous system myelin is required to preserve the integrity of myelin and axons. Glia 67, 634–649 (2019).

16 Yeung, M. S. et al. Dynamics of oligodendrocyte generation and myelination in the human brain. Cell 159, 766–774 (2014).

17 Ringman, J. M. et al. Diffusion tensor imaging in preclinical and presymptomatic carriers of familial Alzheimer's disease mutations. Brain 130, 1767–1776 (2007).

18 Stricker, N. H. et al. Decreased white matter integrity in late-myelinating fiber pathways in Alzheimer's disease supports retrogenesis. Neuroimage 45, 10–16 (2009).

19 Dean, D. C. et al. Association of amyloid pathology with myelin alteration in preclinical Alzheimer disease. JAMA neurology 74, 41–49 (2017).

20 Wang, Q. et al. Quantification of white matter cellularity and damage in preclinical and early symptomatic Alzheimer's disease. NeuroImage: Clinical 22, 101767 (2019).

21 Araque Caballero, M. Á. et al. White matter diffusion alterations precede symptom onset in autosomal dominant Alzheimer’s disease. Brain 141, 3065–3080 (2018).

22 Snaidero, N. et al. Antagonistic functions of MBP and CNP establish cytosolic channels in CNS myelin. Cell reports 18, 314–323 (2017).

23 Lappe-Siefke, C. et al. Disruption of Cnp1 uncouples oligodendroglial functions in axonal support and myelination. Nature genetics 33, 366–374 (2003).

24 Klugmann, M. et al. Assembly of CNS myelin in the absence of proteolipid protein. Neuron 18, 59–70 (1997).

25 Trevisiol, A. et al. Structural myelin defects are associated with low axonal ATP levels but rapid recovery from energy deprivation in a mouse model of spastic paraplegia. PLoS biology 18, e3000943 (2020).

26 Renier, N. et al. iDISCO: a simple, rapid method to immunolabel large tissue samples for volume imaging. Cell 159, 896–910 (2014).

27 Liebmann, T. et al. Three-dimensional study of Alzheimer’s disease hallmarks using the iDISCO clearing method. Cell reports 16, 1138–1152 (2016).

28 Keszycki, R. M., Fisher, D. W. & Dong, H. The hyperactivity–impulsivity– irritiability–disinhibition–aggression–agitation domain in Alzheimer’s disease: current management and future directions. Frontiers in pharmacology 10, 1109 (2019).

29 Chung, J. A. & Cummings, J. L. Neurobehavioral and neuropsychiatric symptoms in Alzheimer's disease: characteristics and treatment. Neurologic clinics 18, 829–846 (2000).

30 Cherny, R. A. et al. Treatment with a copper-zinc chelator markedly and rapidly inhibits β-amyloid accumulation in Alzheimer's disease transgenic mice. Neuron 30, 665–676 (2001).

31 Frenkel, D., Maron, R., Burt, D. S. & Weiner, H. L. Nasal vaccination with a proteosome-based adjuvant and glatiramer acetate clears β-amyloid in a mouse model of Alzheimer disease. The Journal of clinical investigation 115, 2423–2433 (2005).

32 Herculano-Houzel, S., Manger, P. R. & Kaas, J. H. Brain scaling in mammalian evolution as a consequence of concerted and mosaic changes in numbers of neurons and average neuronal cell size. Frontiers in neuroanatomy 8, 77 (2014).

33 Schoenemann, P. T., Sheehan, M. J. & Glotzer, L. D. Prefrontal white matter volume is disproportionately larger in humans than in other primates. Nature neuroscience 8, 242–252 (2005).

34 Stokin, G. B. et al. Axonopathy and transport deficits early in the pathogenesis of Alzheimer's disease. Science 307, 1282–1288 (2005).

35 Gowrishankar, S., Wu, Y. & Ferguson, S. M. Impaired JIP3-dependent axonal lysosome transport promotes amyloid plaque pathology. Journal of Cell Biology 216, 3291–3305 (2017).

36 Vagnoni, A. et al. Calsyntenin-1 mediates axonal transport of the amyloid precursor protein and regulates Aβ production. Human molecular genetics 21, 2845–2854 (2012).

37 Ye, X. & Cai, Q. Snapin-mediated BACE1 retrograde transport is essential for its degradation in lysosomes and regulation of APP processing in neurons. Cell reports 6, 24–31 (2014).

38 Niederst, E. D., Reyna, S. M. & Goldstein, L. S. Axonal amyloid precursor protein and its fragments undergo somatodendritic endocytosis and processing. Molecular biology of the cell 26, 205–217 (2015).

39 Buxbaum, J. D. et al. Alzheimer amyloid protein precursor in the rat hippocampus: transport and processing through the perforant path. Journal of Neuroscience 18, 9629–9637 (1998).

40 Lazarov, O., Lee, M., Peterson, D. A. & Sisodia, S. S. Evidence that synaptically released β-amyloid accumulates as extracellular deposits in the hippocampus of transgenic mice. Journal of Neuroscience 22, 9785–9793 (2002).

41 Gowrishankar, S. et al. Massive accumulation of luminal protease-deficient axonal lysosomes at Alzheimer’s disease amyloid plaques. Proceedings of the National Academy of Sciences 112, E3699–E3708 (2015).

42 Sadleir, K. R. et al. Presynaptic dystrophic neurites surrounding amyloid plaques are sites of microtubule disruption, BACE1 elevation, and increased Aβ generation in Alzheimer’s disease. Acta neuropathologica 132, 235–256 (2016).

43 Yuan, P. et al. TREM2 haplodeficiency in mice and humans impairs the microglia barrier function leading to decreased amyloid compaction and severe axonal dystrophy. Neuron 90, 724–739 (2016).

44 Zhou, Y. et al. Human and mouse single-nucleus transcriptomics reveal TREM2-dependent and TREM2-independent cellular responses in Alzheimer’s disease. Nature medicine 26, 131–142 (2020).

45 Parhizkar, S. et al. Loss of TREM2 function increases amyloid seeding but reduces plaque-associated ApoE. Nature neuroscience 22, 191–204 (2019).

46 Keren-Shaul, H. et al. A unique microglia type associated with restricting development of Alzheimer’s disease. Cell 169, 1276–1290. e1217 (2017).

47 Krasemann, S. et al. The TREM2-APOE pathway drives the transcriptional phenotype of dysfunctional microglia in neurodegenerative diseases. Immunity 47, 566–581. e569 (2017).

48 Huynh, T.-P. V., Davis, A. A., Ulrich, J. D. & Holtzman, D. M. Apolipoprotein E and Alzheimer’s disease: the influence of apolipoprotein E on amyloid-β and other amyloidogenic proteins: Thematic Review Series: ApoE and Lipid Homeostasis in Alzheimer's Disease. Journal of lipid research 58, 824–836 (2017).

49 Hammond, T. R. et al. Single-cell RNA sequencing of microglia throughout the mouse lifespan and in the injured brain reveals complex cell-state changes. Immunity 50, 253–271. e256 (2019).

50 Deming, Y. et al. The MS4A gene cluster is a key modulator of soluble TREM2 and Alzheimer’s disease risk. Science translational medicine 11 (2019).

51 Schirmer, L. et al. Neuronal vulnerability and multilineage diversity in multiple sclerosis. Nature 573, 75–82 (2019).

52 Bartzokis, G. Age-related myelin breakdown: a developmental model of cognitive decline and Alzheimer’s disease. Neurobiology of aging 25, 5–18 (2004).

53 Heneka, M. T. et al. Neuroinflammation in Alzheimer’s disease. The Lancet Neurology 14, 388–405 (2015).

54 Braak, H. & Braak, E. Development of Alzheimer-related neurofibrillary changes in the neocortex inversely recapitulates cortical myelogenesis. Acta neuropathologica 92, 197–201 (1996).

55 C Schmued, L., Raymick, J., G Paule, M., Dumas, M. & Sarkar, S. Characterization of myelin pathology in the hippocampal complex of a transgenic mouse model of Alzheimer’s disease. Current Alzheimer Research 10, 30–37 (2013).

56 Desai, M. K. et al. Early oligodendrocyte/myelin pathology in Alzheimer’s disease mice constitutes a novel therapeutic target. The American journal of pathology 177, 1422–1435 (2010).

57 Desai, M. K. et al. Triple‐transgenic Alzheimer’s disease mice exhibit region‐specific abnormalities in brain myelination patterns prior to appearance of amyloid and tau pathology. Glia 57, 54–65 (2009).

58 Mitew, S. et al. Focal demyelination in Alzheimer’s disease and transgenic mouse models. Acta neuropathologica 119, 567–577 (2010).

59 Roher, A. E. et al. Increased Aβ peptides and reduced cholesterol and myelin proteins characterize white matter degeneration in Alzheimer’s disease. Biochemistry 41, 11080–11090 (2002).

60 Chen, J.-F. et al. Enhancing myelin renewal reverses cognitive dysfunction in a murine model of Alzheimer’s disease. Neuron (2021).

61 Mathys, H. et al. Single-cell transcriptomic analysis of Alzheimer’s disease. Nature 570, 332–337 (2019).

## Methods References

62 Oakley, H. et al. Intraneuronal β-amyloid aggregates, neurodegeneration, and neuron loss in transgenic mice with five familial Alzheimer’s disease mutations: potential factors in amyloid plaque formation. Journal of Neuroscience 26, 10129–10140 (2006).

63 Saito, T. et al. Single App knock-in mouse models of Alzheimer’s disease. Nature neuroscience 17, 661–663 (2014).

64 Meschkat, M. et al. White matter integrity requires continuous myelin synthesis at the inner tongue. bioRxiv (2020).

65 Gorski, J. A. et al. Cortical excitatory neurons and glia, but not GABAergic neurons, are produced in the Emx1-expressing lineage. Journal of Neuroscience 22, 6309–6314 (2002).

66 Berghoff, S. A. et al. Microglia facilitate repair of demyelinated lesions via post-squalene sterol synthesis. Nature neuroscience 24, 47–60 (2021).

67 Singmann, H. et al. afex: analysis of factorial experiments. R package version 0.16-1. R Package Version 0.16 1 (2016).

68 Youmans, K. L. et al. Intraneuronal Aβ detection in 5×FAD mice by a new Aβ-specific antibody. Molecular neurodegeneration 7, 1–14 (2012).

69 Wirths, O. et al. N-truncated Aβ 4–x peptides in sporadic Alzheimer’s disease cases and transgenic Alzheimer mouse models. Alzheimer’s research & therapy 9, 1–12 (2017).

70 Schindelin, J. et al. Fiji: an open-source platform for biological-image analysis. Nature methods 9, 676–682 (2012).

71 Weil, M.-T., Ruhwedel, T., Meschkat, M., Sadowski, B. & Möbius, W. in Oligodendrocytes: Methods and protocols 343–375 (2019).

72 Dobin, A. et al. STAR: ultrafast universal RNA-seq aligner. Bioinformatics 29, 15–21 (2013).

73 Liao, Y., Smyth, G. K. & Shi, W. featureCounts: an efficient general purpose program for assigning sequence reads to genomic features. Bioinformatics 30, 923–930 (2014).

74 Love, M. I., Huber, W. & Anders, S. Moderated estimation of fold change and dispersion for RNA-seq data with DESeq2. Genome biology 15, 1–21 (2014).

75 Corces, M. R. et al. An improved ATAC-seq protocol reduces background and enables interrogation of frozen tissues. Nature methods 14, 959–962 (2017).

76 Stuart, T. et al. Comprehensive integration of single-cell data. Cell 177, 1888–1902. e1821 (2019).

77 McInnes, L., Healy, J. & Melville, J. Umap: Uniform manifold approximation and projection for dimension reduction. arXiv preprint arXiv:1802.03426 (2018).

78 Finak, G. et al. MAST: a flexible statistical framework for assessing transcriptional changes and characterizing heterogeneity in single-cell RNA sequencing data. Genome biology 16, 1–13 (2015).

79 Barrett, T. et al. NCBI GEO: archive for functional genomics data sets— update. Nucleic acids research 41, D991–D995 (2012).

